# Antibacterial Compounds Against Non-Growing and Intracellular Bacteria

**DOI:** 10.1101/2024.11.06.622235

**Authors:** Niilo Kaldalu, Normunds Bērziņš, Stina Berglund Fick, Atin Sharma, Naomi Charlotta Andersson, Jüri Aedla, Mariliis Hinnu, Andrea Puhar, Vasili Hauryliuk, Tanel Tenson

## Abstract

Slow- and non-growing bacterial populations, along with intracellular pathogens, often evade standard antibacterial treatments and are linked to persistent and recurrent infections. This necessitates the development of therapies specifically targeting nonproliferating bacteria. To identify compounds active against non-growing uropathogenic *Escherichia coli* (UPEC) we performed a drug-repurposing screen of 6,454 approved drugs and drug candidates. Using dilution-regrowth assays, we identified 39 compounds that either kill non-growing UPEC or delay its regrowth post-treatment. The hits include fluoroquinolones, macrolides, rifamycins, biguanide disinfectants, a pleuromutilin, and anti-cancer agents. 29 of the hits have not previously been recognized as active against non-growing bacteria. The hits were further tested against non-growing *Pseudomonas aeruginosa* and *Staphylococcus aureus*. Ten compounds – solithromycin, rifabutin, mitomycin C, and seven fluoroquinolones – have strong bactericidal activity against non-growing *P. aeruginosa*, killing >4 log_10_ of bacteria at 2.5 µM. Solithromycin, valnemulin, evofosfamide, and satraplatin are unique in their ability to selectively target non-growing bacteria, exhibiting poor efficacy against growing bacteria. Finally, 31 hit compounds inhibit the growth of intracellular *Shigella flexneri* in a human enterocyte infection model, indicating their ability to permeate the cytoplasm of host cells. The identified compounds hold potential for treating persistent infections, warranting further comparative studies with current standard-of-care antibiotics.

## INTRODUCTION

All currently used antibiotics were discovered based on their ability to inhibit bacterial growth^1^. However, only a few can effectively kill non-growing bacteria, which are prevalent in chronic and recurrent infections^2^. The ability of bacteria to survive treatment with bactericidal antibiotics is termed antibiotic tolerance, distinct from antibiotic resistance, which refers to the ability to grow in the presence of the drug^3^. Non-growing bacteria, being tolerant to antibiotics, can evade conventional antibacterial therapy and are more likely to develop resistance^4,5^. Bacterial cultures commonly contain a subpopulation of non-growing cells, known as persisters, that are tolerant to bactericidal antibiotics^6–8^. However, the connection between these *in vitro* persisters and persistent infections in the clinical settings remains uncertain and is challenging to investigate^9,10^. Furthermore, chronic infections are often caused by intracellular bacteria as well as by biofilms, which both evade the immune defense and antibiotics^11–14^. To effectively treat these infections and curb the spread of resistance, there is an urgent need for drugs that selectively target non-growing and intracellular bacteria. These drugs must be devoid of severe side effects and either possess antibacterial activity on their own or enhance conventional antibiotics.

In this study, we performed a high-throughput screening to identify compounds that selectively target non-growing Gram-negative bacteria using a dilution-regrowth assay^15^. Specifically, drug-treated stationary phase culture samples were diluted into fresh medium, and the time to outgrowth was measured as a proxy for the number of bacteria that survived the treatment. Using a library of approved and candidate drugs with known safety and pharmacokinetic properties excluded reactive and toxic compounds that might otherwise come up as false hits in the screen.

In our primary screen we used uropathogenic *Escherichia coli* (UPEC), a clinically important pathogen and a model organism for other Gram-negative pathogens. UPEC is a prevalent cause of recurrent urinary tract infections (UTIs)^16^, which are among the most common outpatient bacterial infections^17^ and exhibit high treatment failure rates^18,19^. UPEC can reside both extracellularly and intracellularly, forming persistent intracellular bacterial reservoirs in urinary epithelial cells and macrophages^20,21^. Reactivation of quiescent bacteria in the urinary tract may be one cause of recurrent UTIs, along with potential reinfection originating from the host’s intestinal tract^22–24^. To identify drug candidates targeting intracellularly persisting bacteria, the screen was performed in an acidic, low-phosphate, and low-magnesium medium that is designed to mimic conditions in intravacuolar reservoirs, where quiescent UPEC resides within host cells^22,25^.

After characterizing the effects of the hit compounds on non-growing UPEC, we tested their activities against stationary phase cultures of *Pseudomonas aeruginosa* and *Staphylococcus aureus* to explore their potential broader use. These two pathogens were selected due to their prevalence in chronic biofilm infections^26,27^. Finally, to assess the capacity of the hit compounds to permeate through the host cell cytoplasmic membrane barrier, we characterized their inhibitory activity against *Shigella flexneri* replicating in the cytosol of intestinal epithelial cells^28^.

## RESULTS

### A dilution-regrowth screen identifies compounds that delay the regrowth of non-growing UPEC

We conducted a screen of 6,454 registered drugs and drug candidates, aiming to identify compounds effective against non-growing UPEC strain CFT073^29^. This collection comprised 1,200 compounds from the Prestwick Library and 5,254 from the Specs Repurposing Library. The Prestwick Library primarily consists of off-patent, regulator-approved drugs from diverse therapeutic classes, including 144 antibiotics and 12 antiseptics, while the Specs Library contains commercially available drugs, drug candidates in clinical development phases 1 through 3, and compounds in preclinical development. All compounds were tested at a final concentration of 20 μM.

Bacteria were cultured in either 1:4 diluted cation-adjusted Mueller-Hinton broth (CA-MHB) at pH 7.4 or in an acidic, low-phosphate, low-magnesium medium (LPM) at pH 5.5. CA-MHB was used diluted to reduce bacterial clumping in dense stationary-phase cultures and thereby decrease pipetting error. Acidic LPM was chosen to simulate conditions in UPEC-inhabited vacuoles^22,25^. Given that variation in antibiotic tolerance decreases when cultures remain in the stationary phase for extended periods or are inoculated from such cultures^30,31^, we used a 24- hour cultivation and treatment time in both the screening and validation experiments to minimize stochasticity and variability.

The screening assay involved treating a stationary-phase culture with each compound for 24 h, followed by monitoring the delay in bacterial outgrowth after a 2,500-fold dilution into drug-free growth medium (Fig. 1A)^15^. This dilution reduced the compound concentration to 8 nM, which was expected to be below their minimum inhibitory concentration (MIC), thus allowing the regrowth of surviving bacteria. This assumption was confirmed, as the lowest measured MIC for all hit compounds was 12.5 nM (clinafloxacin, see Dataset 1).

**Figure 1.**
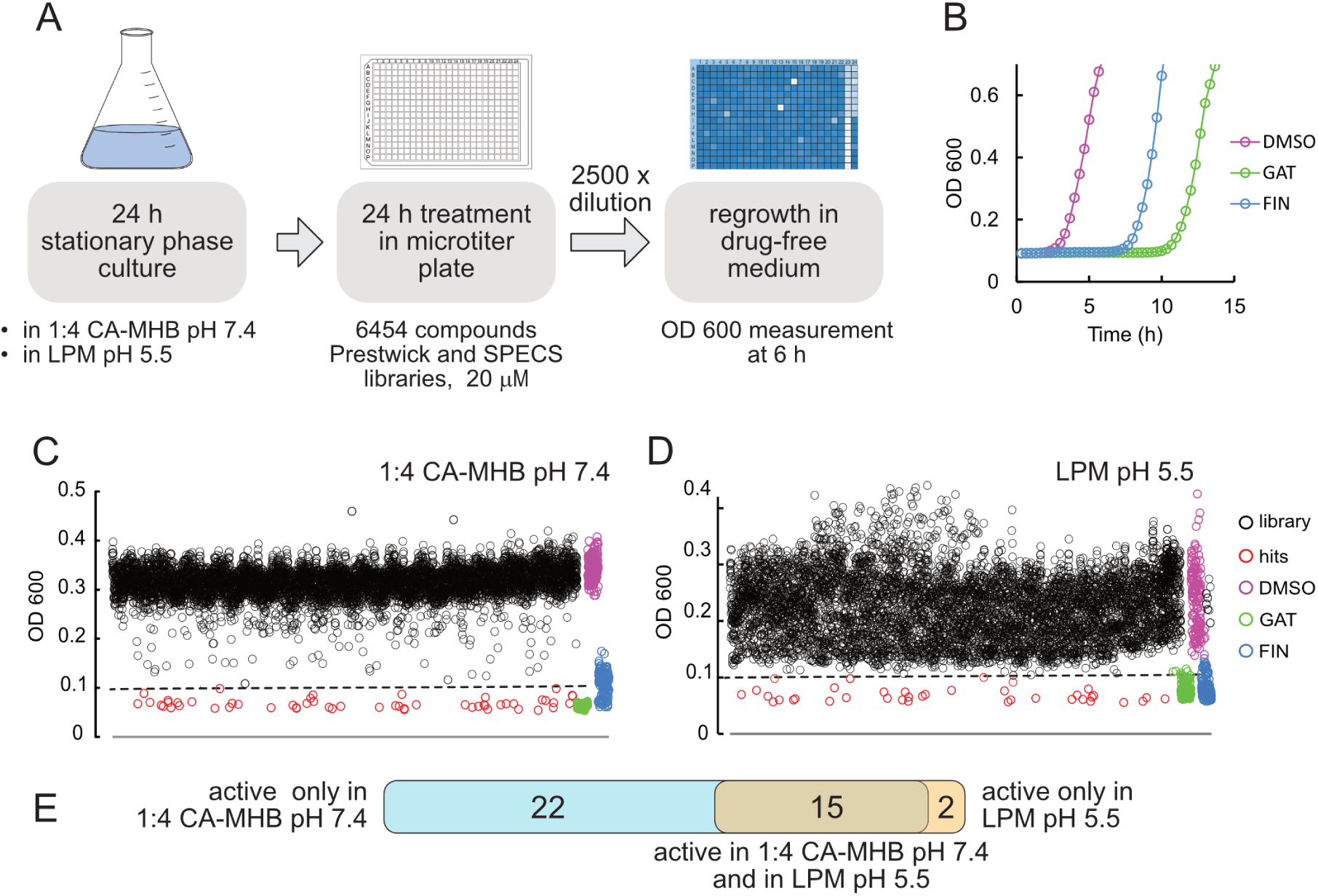
A dilution-regrowth-based screen identifies compounds that delay the regrowth of stationary-phase UPEC. **A.** Schematic of the screening assay. A 24-h culture of UPEC strain CFT073 was dispensed into a microtiter plate and treated with 20 μM compounds from the collection for 24 h. The samples were then diluted 2,500-fold into drug-free CA-MHB (pH 7.4), and OD_600_ was measured after 6 h. **B.** Regrowth delay following bactericidal treatment. For assay validation, UPEC CFT073 was cultivated for 24 h in 1:4 diluted CA-MHB (pH 7.4) and treated for 24 h with 20 μM gatifloxacin (GAT) or finafloxacin (FIN), which kill non-growing bacteria and served as positive controls for screening. After a 2,500-fold dilution into drug-free CA-MHB, the regrowth of the drug-treated samples was delayed compared to that of the drug-free control (DMSO). **C, D.** Screening of the combined Prestwick and SPECS collections identified drugs that delayed bacterial regrowth in 1:4 diluted CA-MHB (pH 7.4) (C) and LPM (pH 5.5) (D). A cutoff of OD_600_ < 0.1 (dashed line) was used to identify hits (red). Most of the tested compounds (black) showed no effect compared to the drug-free control (purple). GAT (green) and FIN (blue) were included as positive controls in each plate. **E**. A diagram showing validated hits from either 1:4 CA-MHB (pH 7.4), LPM (pH 5.5), or both media.

To validate the assay, we used gatifloxacin (GAT) and finafloxacin (FIN) as positive controls, while mock-treated samples (1% DMSO without antibiotics) served as negative controls (Fig. 1B, Fig. S1). GAT has previously been shown to kill UPEC persisters ^32^, while FIN has bactericidal activity at acidic pH and kills effectively intracellular UPEC in a cell culture model^33,34^. Preliminary experiments confirm that both GAT and FIN have bactericidal effects against stationary-phase UPEC cultures at a concentration of 20 µM. The Z’-factor, indicating assay robustness and discrimination between positive controls and mock treatments, was above 0.5 between 5 and 8 h after dilution (Fig. S1), demonstrating excellent assay reliability within this timespan.

In the course of the screening, we measured the optical density of regrowing samples 6 h post-dilution, defining wells with an OD_600_ < 0.1 as hits. GAT, FIN, and DMSO controls were included on every screening plate to ensure consistency (Fig. 1C,D). After eliminating duplicates — compounds present in both libraries or those appearing in different chemical forms (e.g., the hydrochloride salt and free base of the same fluoroquinolone) — we identified 54 hits in 1:4 CA-MHB pH 7.4 and 38 hits in LPM pH 5.5, with 23 overlapping hits active in both media.

### Hit validation and dose-response analysis

Hits from the primary screen were validated through a dilution-regrowth assay at 20, 10, and 5 μM concentrations in both 1:4 CA-MHB pH 7.4 and LPM pH 5.5 (Fig. S2). Compounds that delayed regrowth, resulting in an OD_600_ at least 50% lower than the drug-free control at one or more of these concentrations after 6 h of incubation in regrowth medium, were selected for further testing at concentrations ranging from 30 to 0.25 μM. We identified 39 compounds of six different classes that significantly delayed regrowth at one or more of these concentrations. Of these compounds, 37 were active in 1:4 CA-MHB, 17 in acidic LPM and 15 in both media (Fig. 1E, 2, and 3).

**Figure 2.**
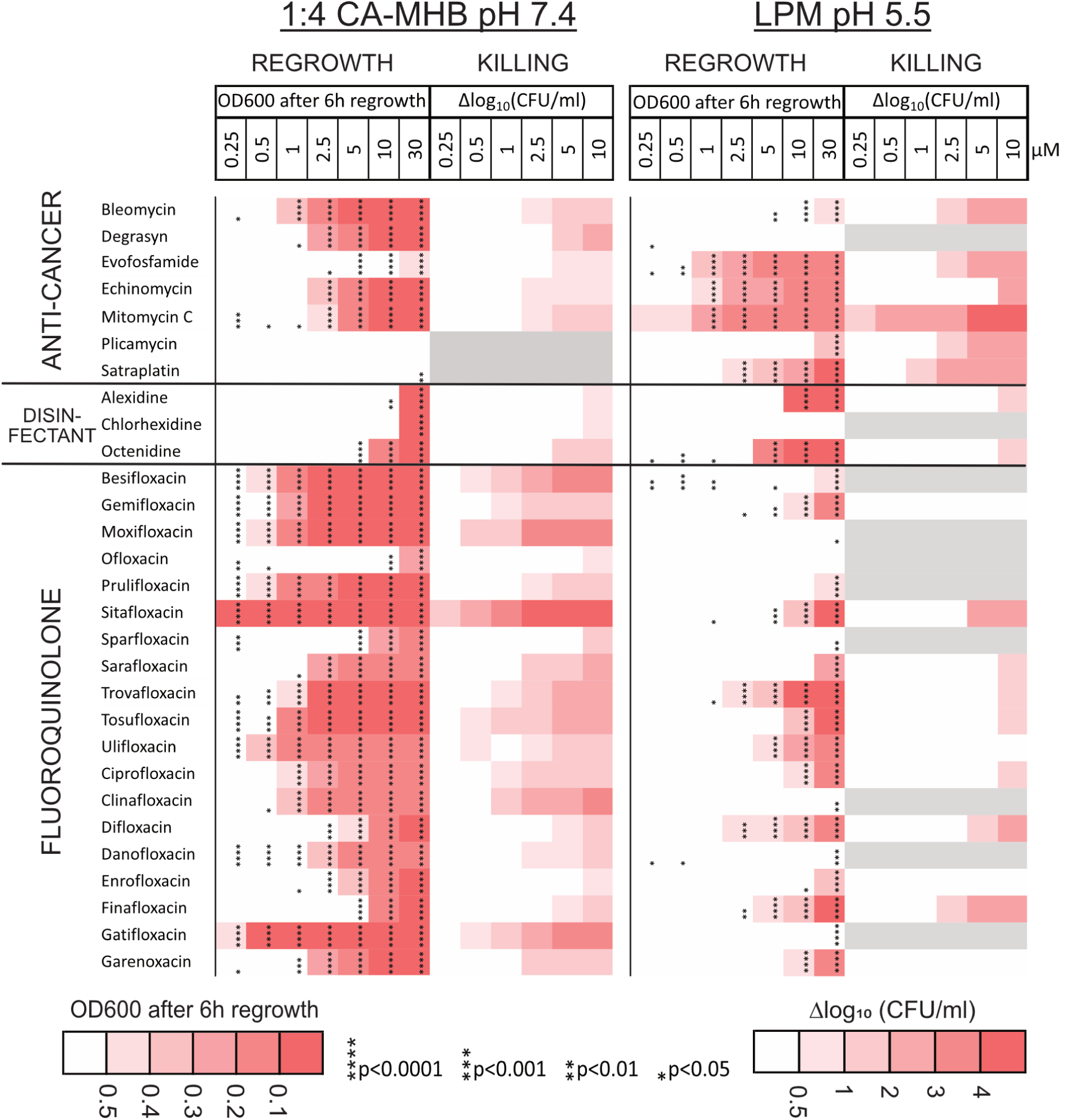
Regrowth delay and killing of non-growing UPEC. Heatmap showing the regrowth delay and killing of non-growing UPEC CFT073 by validated hit compounds. Gray areas indicate compounds not tested for killing due to weak activity in regrowth-delay experiments. Bacteria were grown in 1:4 CA-MHB (pH 7.4) or LPM (pH 5.5) for 24 h and then treated with the compounds at the indicated concentrations for 24 h. Regrowth data were obtained by measuring OD_600_ 6h after a 2,500-fold dilution of the samples into CA- MHB. Killing data were determined by spot-plating serial dilutions of the treated samples on LB agar plates followed by colony counting. The differences in CFU/mL between the drug-treated samples and the drug-free control are shown. Data represent the mean of three biological replicates. The significance of the regrowth delay was assessed using one-way ANOVA followed by Dunnett’s test, comparing each treatment to the drug-free control. Asterisks denote p-values from the Dunnett’s test.

**Figure 3.**
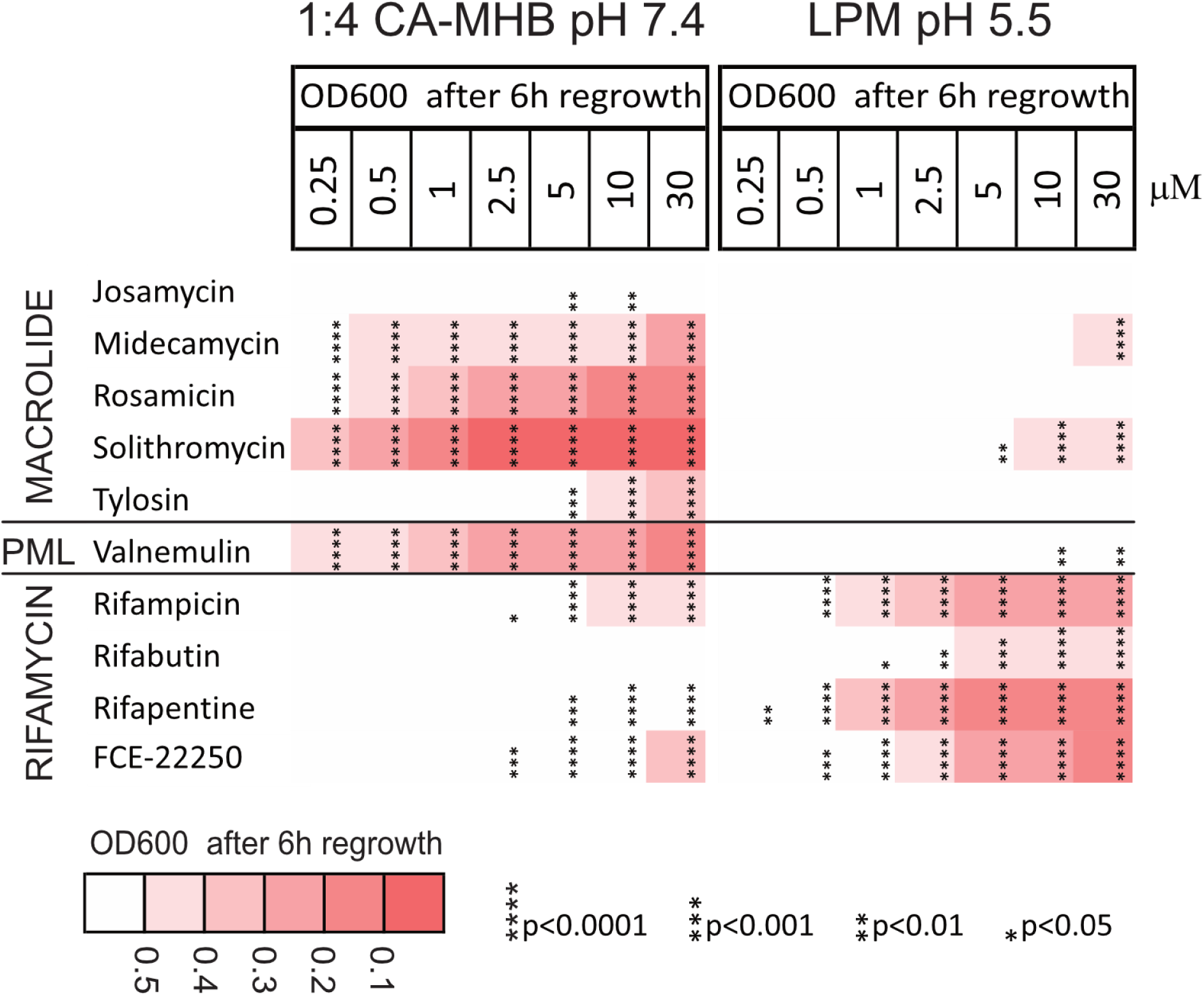
The post-antibiotic effect on non-growing UPEC. Heatmap showing the regrowth delay of non-growing UPEC CFT073 by validated hit compounds that did not kill the stationary phase bacteria. Cultures were grown in 1:4 CA-MHB (pH 7.4) or LPM (pH 5.5) for 24 h and then treated with the compounds at the indicated concentrations for another 24 h. OD_600_ was measured 6 h after a 2,500-fold dilution in CA- MHB. PML refers to pleuromutilin. Data represent the average of three biological replicates. Regrowth delay significance was assessed using one-way ANOVA and Dunnett’s test, with comparisons made against the drug-free control. Asterisks indicate p-values from Dunnett’s test.

Of the 39 compounds, 29 are well-known antibiotics. While ofloxacin, ciprofloxacin^35,36^, tosufloxacin, sparfloxacin, moxifloxacin, gatifloxacin, enrofloxacin, sarafloxacin^32,37^ as well as rifampicin ^38,39^ have been previously identified as active against non-growing bacteria, to the best of our knowledge, this has not been demonstrated for the remaining 20 antibiotics that were hits from the screen. The largest group consists of 19 fluoroquinolones, including 18 hits from the screen, as well as finafloxacin which was used as a positive control. Other antibiotic classes include protein synthesis inhibitors: five macrolides and one pleuromutilin (valnemulin), as well as four RNA synthesis-inhibiting rifamycins. The list also includes three antiseptics and disinfectants: the biguanide octenidine, and the bisbiguanides alexidine and chlorhexidine. The antibacterial activity of biguanides and bisbiguanides is attributed to their membrane-disrupting activity^40^. In contrast, colistin, a membrane-targeting last-resort antibiotic, was not among the hits.

Hit compounds that are neither antibiotics nor cationic surfactants, are anti-cancer agents with varying mechanisms of action and at different stages of development. Mitomycin C (MMC)^41^, bleomycin (BLE)^42^, satraplatin (SAT)^43^, and evofosfamide (EVO)^44^ are known genotoxic compounds targeting DNA integrity and synthesis. Echinomycin and plicamycin (also known as mithramycin) are DNA-intercalating antitumor drugs that block transcription^45^. A deubiquitinase inhibitor Degrasyn (DGS, WP1130) is a thiol-reactive compound that modifies cysteine residues^46,47^.

In summary, our screen identified numerous compounds that were previously unknown to be active against non-growing UPEC.

### Both killing and the post-antibiotic effect contribute to delaying UPEC regrowth

While setting up the screen, we anticipated that the killing of non-growing bacteria would be the primary cause of the regrowth delay. An alternative cause could be the post-antibiotic effect (PAE) without killing, i.e., a delay in the post-treatment regrowth of individual bacteria after an antibiotic is removed from the extracellular environment^48^. It is also possible that both bacterial killing and PAE occur simultaneously.

To discriminate between these two scenarios, we assessed the bactericidal effect on non-growing bacteria by measuring the reduction in colony-forming units (CFU) by agar plating for all compounds that demonstrated activity in the dilution-regrowth assay at a concentration of 10 μM (Fig. S2). Fluoroquinolones, anti-cancer agents, and disinfectants do kill a fraction of stationary phase UPEC (Fig. 2), whereas macrolides, rifamycins, and valnemulin are not bactericidal against *E. coli* but rather cause PAE (Fig. 3, Dataset 2). The occurrence of PAE indicates that these drugs penetrate the non-growing bacteria and remain there for hours, inhibiting targets and postponing regrowth. Fluoroquinolones are known to cause PAE after treatment of growing bacterial cultures^49^. We compared the bactericidal activity of each compound based on the log killing at a 2.5 μM concentration and regrowth-inhibiting activity by determining the EC_50_ values, which were calculated using the OD_600_ readings from treated and diluted stationary phase samples after a 6 h incubation period. Here, EC_50_ represents the concentration of a compound that causes a regrowth delay, halfway between the baseline (no regrowth inhibition, with OD_600_ of the drug-free control) and the maximum effect (complete inhibition of regrowth, with OD_600_ of the sterile medium) (Dataset 1). We found that fluoroquinolones with similar EC_50_ exhibit different bactericidal activity, although these two factors are correlated (r = -0.67; 95% CI = -0.86 to -0.29; p = 0.0025) (Fig. S3A). This indicates that both reduction of the inoculum by killing and the PAE of the surviving bacteria contributed to the observed regrowth delay by fluoroquinolones (Fig. 2). Finally, we found that sitafloxacin is the most potent and bactericidal compound against non-growing UPEC at pH7.4, considerably exceeding ciprofloxacin, which is recommended for the treatment of uncomplicated pyelonephritis and complicated UTIs^50,51^.

### The activity against non-growing UPEC depends on the pH of the medium

The majority of the hit compounds were less effective in LPM pH5.5 compared to 1:4 CA- MHB pH7.4. Several fluoroquinolones, macrolides, valnemulin, and chlorhexidine were active at pH7.4 and completely inactive in the acidic medium (Fig. 2, 3). In particular, the bactericidal activity of all the fluoroquinolones, except finafloxacin, drops significantly at acidic pH (Fig. 2, Fig. S3A,B). Conversely, the rifamycins, alexidine, and octenidine, as well as the anti-cancer agents evofosfamide, mitomycin C, satraplatin, and plicamycin, are potentiated by the low pH of the medium (Fig. 2, 3). In total, eleven compounds are more active in LPM pH5.5 than in 1:4 CA-MHB pH7.4 (Fig. S4).

### Hit compounds have different activity profiles against non-growing P. aeruginosa and S. aureus

To determine the spectrum of activity of the compounds that were effective against non-growing UPEC, we further assessed their efficacy against the well-characterized Gram-negative and Gram-positive reference strains *P. aeruginosa* DSM1117 and *S. aureus* DSM2569, respectively^52^. The cultures were grown and treated for 24 h in 1:4 CA-MHB pH7.4. 24 compounds that were identified as the most active in preliminary dilution-regrowth tests at a concentration of 10 μM were further characterized for regrowth delay and bactericidal activity at concentrations ranging from 10 to 0.1 μM (Fig. S2).

For better characterization of the compounds causing the strongest regrowth delay, we performed the regrowth step of the dilution-regrowth experiments on a plate reader, measuring the OD_600_ at 20-minute intervals for over 20 h instead of using an endpoint measurement. This approach allowed us to capture the time when cultures reached a threshold optical density (Start-Growth-Time, SGT^15^).

*P. aeruginosa* is notorious for causing resilient infections that do not respond well to antibiotics^26^. Therefore, we were surprised to find that many drugs, especially fluoroquinolones, were active against non-growing *P. aeruginosa* DSM1117 (Fig. 4). Strikingly, the bactericidal activity of the tested drugs against this *P. aeruginosa* strain is more potent compared to their activity against UPEC CFT073 (Fig. 2,4, Fig. S3A,C). Of the 24 compounds that are most active against *P. aeruginosa* based on preliminary regrowth-dilution results, all are also effective at killing nongrowing bacteria of this species, including those that induced only a post-antibiotic effect (PAE) without reducing CFU in UPEC.

**Figure 4.**
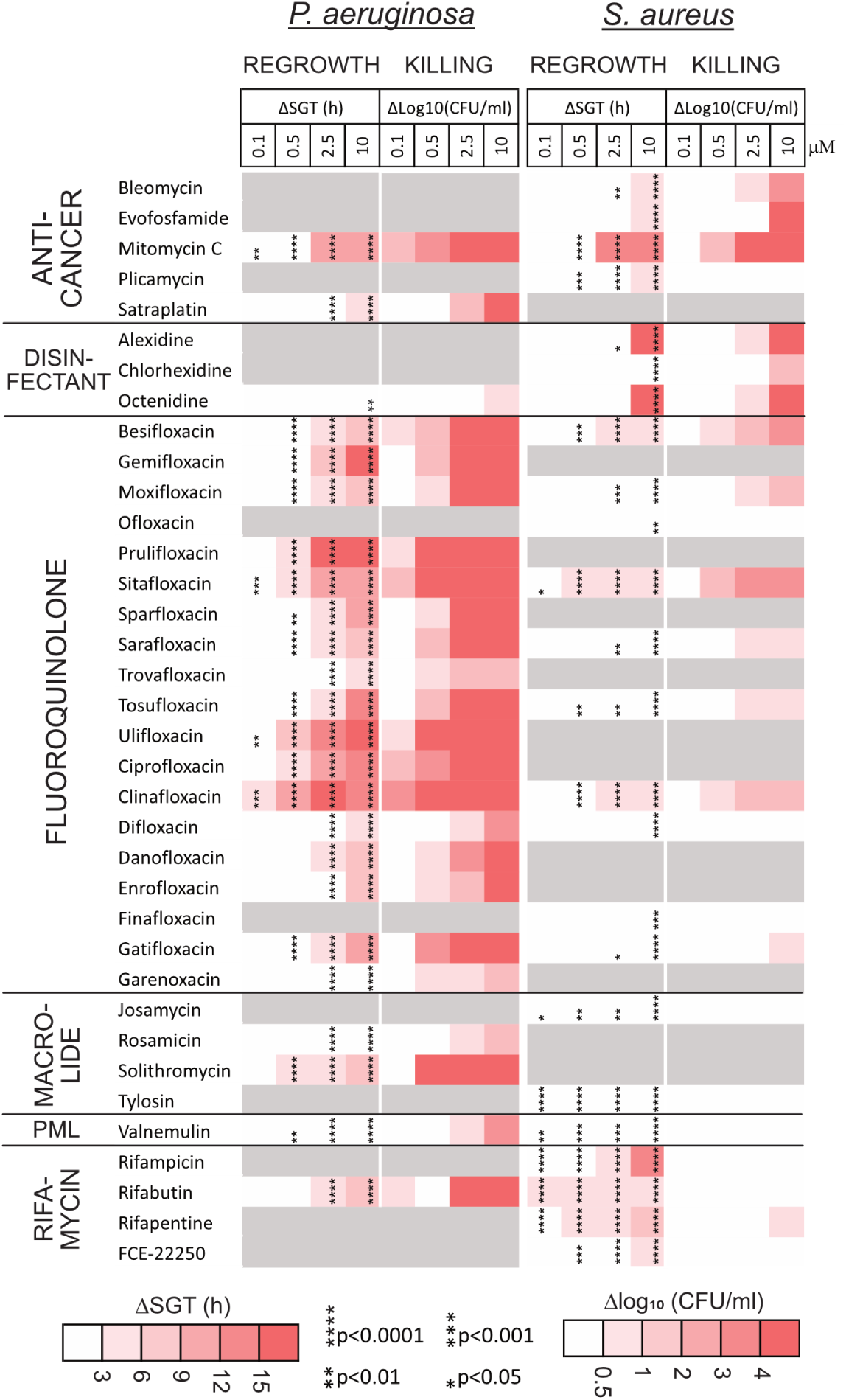
Regrowth delay and killing of non-growing *P. aeruginosa* and *S. aureus*. Regrowth delay and killing of non-growing *P. aeruginosa* DSM1117 and *S. aureus* DSM2569 by compounds effective against non-growing UPEC. Gray areas in the heat map indicate compounds not tested due to inactivity in preliminary experiments. Cultures were grown in 1:4 CA-MHB (pH 7.4) for 24 h followed by treatment with the compounds at the indicated concentrations for an additional 24 h. To assess regrowth, OD_600_ was measured on a microplate reader after a 2,500-fold dilution into CA-MHB. The time required for cultures to reach an OD_600_ threshold of 0.12 is reported as the Start-Growth-Time (SGT). Bacterial killing was determined by spot-plating serial dilutions of the treated samples on LB agar plates followed by colony counting. The differences in CFU/mL between drug-treated samples and the drug-free control are presented. Data represent the average of three biological replicates. The significance of regrowth delay was evaluated using OD_600_ measurements taken 8 h after dilution into drug-free media. Statistical significance was assessed using one-way ANOVA followed by Dunnett’s test, comparing each treatment to the drug-free control. Asterisks denote p-values from Dunnett’s test.

Notably, solithromycin (a macrolide) and rifabutin (a rifamycin) were highly bactericidal against non-growing *P. aeruginosa*, reducing bacterial counts by more than four logs at concentrations of 10 μM and 2.5 μM, respectively. In contrast, these drugs were previously shown not to kill UPEC but only to induce PAE. Solithromycin kills more than four logs at a concentration as low as 0.5 μM, similarly to the most bactericidal fluoroquinolones: clinafloxacin, ulifloxacin, sitafloxacin and prulifloxacin. Altogether, 10 compounds (including the above-listed, as well as mitomycin C, besifloxacin, gemifloxacin, and ciprofloxacin) reduce bacterial counts by more than four logs at a concentration of 2.5 µM. In several individual samples, the bacterial count is reduced to undetectable levels: no colonies were observed on agar from the smallest plated dilution, and/or the diluted bacteria did not regrow in liquid medium during the observation period (Dataset 2).

The activity profile of the hit compounds against non-growing *S. aureus* differed significantly from those against both UPEC and *P. aeruginosa* (Fig. 4, Fig. S3D). In the regrowth-dilution assay, most fluoroquinolones and macrolides are inactive, exhibit weak activity, or show activity only at the highest concentration tested, 10 µM. While macrolides do not kill non-growing bacteria, some of the fluoroquinolones reduce bacterial counts by one to two logs, with sitafloxacin showing the most activity. Rifamycins do not kill non-growing *S. aureus*, but they demonstrate an increased activity in delaying regrowth compared to their effects on the Gram-negative organisms. Together with the disinfectants alexidine and octenidine, the anti-cancer agents mitomycin C and evofosfamide are the most bactericidal compounds in our set, reducing non-growing *S. aureus* below the limit of detection at a concentration of 10 µM (Fig. 4).

### Diverse hit compounds are active against non-growing bacteria at sub-MIC concentrations

The bactericidal effect of antibiotics correlates positively with bacterial growth rate^35,53–55^, implying that slow-growing bacteria are less sensitive to these drugs. Non-growing bacteria are typically tolerant to β-lactams, other cell-wall inhibitors, and aminoglycosides, which did not emerge as hits. In contrast, rapid bacterial growth has been found to reduce macrolide accumulation and efficacy^56^. To characterize the activity of the hit compounds against growing bacteria, we measured their MIC values and found that some had MIC values much higher than the concentrations needed to kill non-growing bacteria or delay their regrowth (Dataset 1).

The MIC of midecamycin against UPEC in a standard assay (in CA-MHB) is over 160 µM, yet it effectively delays UPEC regrowth at 0.5 µM (Fig. 3). Similarly, solithromycin has a MIC of 80 µM against *P. aeruginosa* but kills almost four logs of stationary phase cells at 0.5 µM (Fig. 4). Comparing the activity of hit compounds against growing bacteria (MIC) and non-growing bacteria (EC_50_ for regrowth delay) shows considerable differences between organisms, groups of compounds, and testing conditions (Fig. 5). Anti-cancer agents are inactive based on MIC (EVO, Fig. 5A,B,D; SAT, Fig. 5C) or have an MIC far higher than EC_50_ (MMC in *P. aeruginosa*, Fig. 5C). In Gram-negative *E. coli* and *P. aeruginosa*, the MIC of macrolides and valnemulin considerably exceeds EC_50_ in 1:4 CA-MHB medium (Fig. 5A,C). There was no statistically significant correlation between MIC and EC_50_ for regrowth delay across the entire set of the hit compounds (see Fig. 5 legend for statistical details). For fluoroquinolones, the largest group of compounds, EC_50_ exceeded MIC manyfold for UPEC in 1:4 CA-MHB pH 7.4, indicating that growing *E. coli* is more sensitive to these compounds than non-growing cells (Fig. 5A). For *P. aeruginosa*, however, MIC values are higher than EC_50_ for all fluoroquinolones, suggesting that non-growing bacteria in this strain are more sensitive to these compounds than growing bacteria, which is unexpected (Fig. 5C).

**Figure 5.**
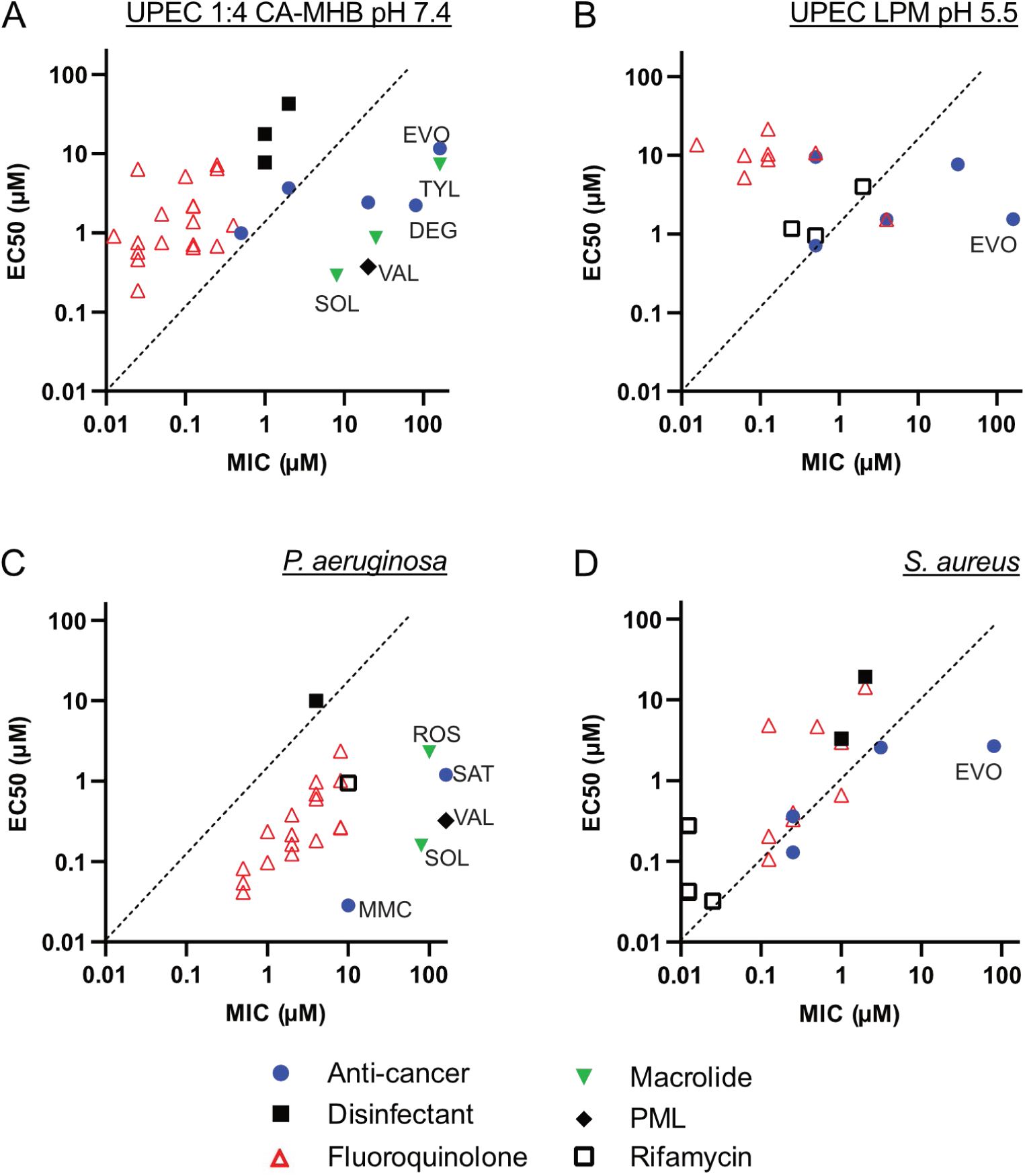
Activity of hit compounds against growing and nongrowing bacteria. The activity against growing bacteria was assessed by determining the minimal inhibitory concentration (MIC) using the broth dilution method. The activity against non-growing bacteria was evaluated by calculating the EC_50_ based on regrowth delay, measured by OD_600_ readings at 6 h (A, B) or 8 h (C, D) after dilution of treated bacteria into drug-free growth media. EVO – evofosfamide; DEG – degrasyn; MMC – mitomycin C; SOL – solithromycin; TYL – tylosin; ROS – rosamicin; VAL – valnemulin. **A, C, D**. 1:4 CA-MHB (pH 7.4) was used for cultivation and treatment of non-*growing* UPEC CFT073, *P. aeruginosa* DSM1117, and *S. aureus* DSM2569. 1:4 CA-MHB (pH 7.4) and CA- MHB (pH 7.4) were used for MIC measurement (Dataset 1). **B**. LPM (pH 5.5) was used in experiments with non-growing *E. coli* CFT073 and MIC determination (Dataset 1).

Comparing UPEC MIC values at pH 7.4 and 5.5, we found that the anti-cancer agents, which were potentiated against non-growing UPEC in acidic LPM, also have lower MIC in this medium. All rifamycins and three fluoroquinolones (finafloxacin, trovafloxacin, and difloxacin) display lower MIC at pH 5.5 compared to pH 7.4 (Dataset 1). For trovafloxacin and difloxacin, this does not translate into superior activity against non-growing UPEC in acidic medium; however, both drugs are active against non-growing UPEC in both media (Fig. 2).

The standard medium for MIC determination is CA-MHB, which we also initially used to determine the MIC of the hit compounds; however, we cultured and treated the stationary-phase bacteria in a diluted medium with a fourfold reduction in divalent cation concentration. The composition of the testing medium significantly impacts both the MIC values and the bactericidal rates^57^. To determine if the disparity between high MIC and activity against non-growing cultures was due to different testing media, we measured the MIC of selected compounds in 1:4 CA-MHB pH 7.4. We found that extremely high MICs generally do not decrease in the diluted medium. Exceptions included two rifamycins: rifabutin (in UPEC and *P. aeruginosa*) and FCE-22250 (in *P. aeruginosa*), whose MICs drop in the diluted medium. Additionally, the MIC of most fluoroquinolones in *P. aeruginosa* remain the same in diluted CA-MHB, or even increase, as in the case of trovafloxacin and besifloxacin (four- and two-fold, respectively; Dataset 1).

Another possible explanation for high activity against non-growing bacteria and lack of activity against growing cells might be the effective pumping out of the compounds from growing bacteria, where efflux pumps are energized and active. To test this, we measured MIC in the presence of the broad-spectrum efflux pump inhibitor phenylalanine-arginine β-naphthylamide (PaβN). PaβN does not reduce the MIC of compounds with the highest MIC, suggesting that efflux does not explain their high MICs (Dataset 1).

### Hit compounds display different kinetics of antimicrobial activity against non-growing bacteria

For screening and concentration-dependent characterization of the hit compounds, we used a 24-hour incubation to capture their effects, although the actual onset of their activity may occur much sooner. To assess the speed of action, we treated stationary-phase cultures of three organisms with compounds that had previously shown a strong regrowth delay at 10 µM. We selected compounds that, in earlier tests, delayed regrowth enough that diluted cultures did not exceed an OD_600_ of 0.12 after spending 6 h for UPEC or 8 h for *P. aeruginosa* and *S. aureus* in drug-free medium. This requirement was met by 18 compounds for *E. coli* CFT073, 21 compounds for *P. aeruginosa*, and 11 compounds for *S. aureus*.

We collected samples at different times during the 24-hour incubation, diluted them 2,500- fold, and measured the OD_600_ after 6 or 8 h of further incubation, respectively. As seen in Fig. 6, most of the compounds cause the growth delay after just 1 h incubation in all three organisms. A few compounds act more slowly, such as garenoxacin in *E. coli*; difloxacin, satraplatin, trovafloxacin, and sparfloxacin in *P. aeruginosa*. Rifampicin acts quickly against non-growing *S. aureus,* while the effects of the other compounds increase gradually during incubation but are already apparent within the first hour of treatment.

**Figure 6.**
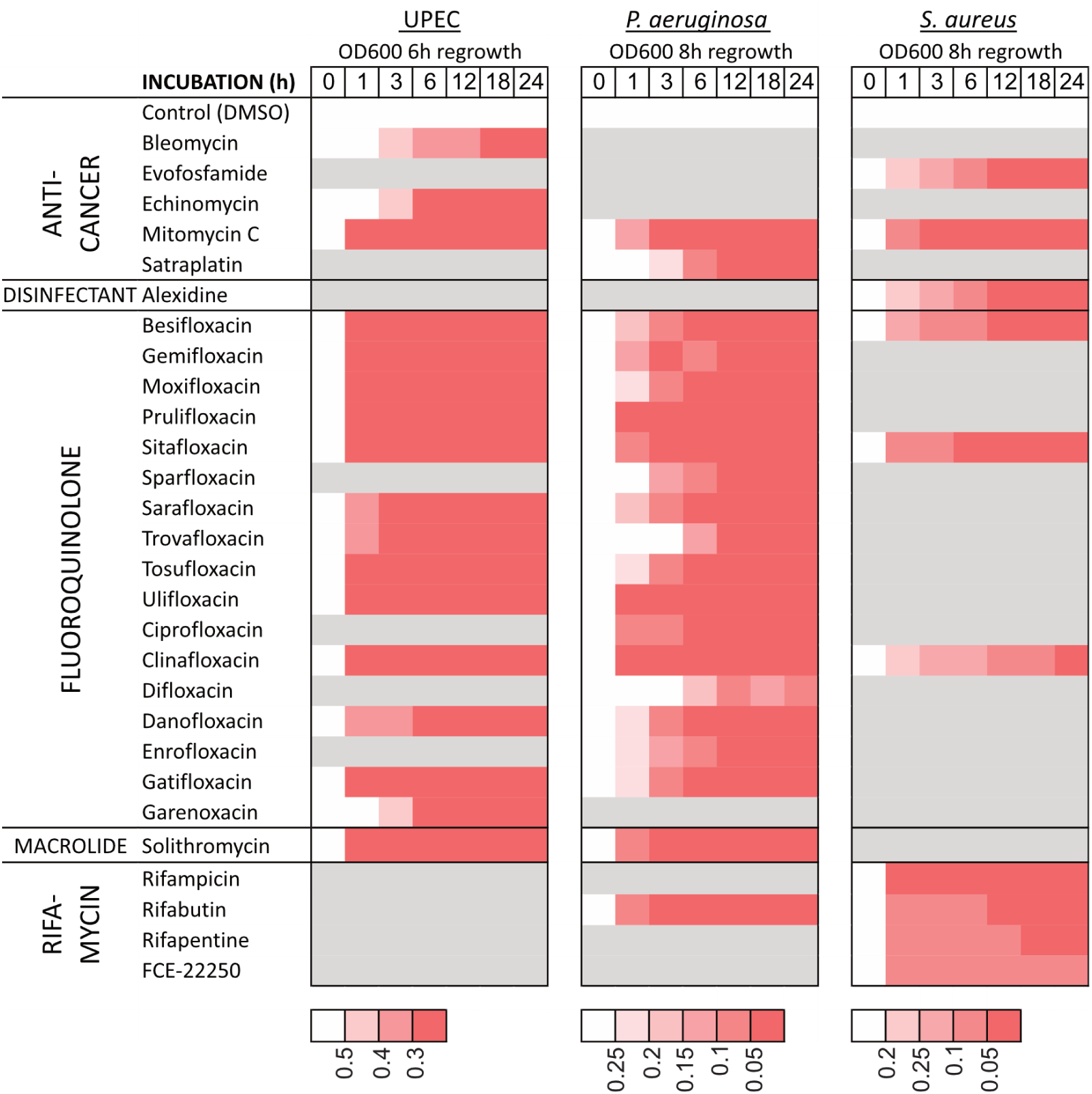
Time course of antibacterial effects on non-growing bacteria. Bacteria were cultivated in 1:4 CA-MHB (pH 7.4) for 24 h and the compounds were added at a concentration of 10 µM. Samples were taken at the indicated time points, diluted 2,500-fold into CA-MHB, and OD_600_ was measured after 6h of incubation for UPEC CFT073 and 8h for *P. aeruginosa* and *S. aureus*. The color scale for OD_600_ values corresponding to each organism is shown below its respective heat map. Gray areas in the heat map indicate compounds not tested due to inactivity observed in previous experiments. Data represent the average of three biological replicates.

In summary, these results indicate that most compounds penetrate non-growing bacteria rapidly, exhibiting an antibacterial effect after one hour of incubation.

### DNA damage is a possible anti-bacterial mechanism of the anti-cancer compounds

Most of the anti-cancer agents (MMC, BLE, SAT and EVO) that exhibit bactericidal activity against non-growing bacteria are genotoxic, which makes their development into antimicrobials unlikely. The other two drugs of this group, echinomycin (ECH) and plicamycin are DNA-intercalating transcription blockers^45^. We aimed to evaluate the genotoxicity of ECH and PLI by detecting the possible induction of the SOS response. Degrasyn was excluded from further testing due to its chemical instability.

The SOS response is induced by DNA damage that leads to double-strand breaks and RecA activation. The SOS regulon consists of several operons that are controlled by the LexA repressor, which is degraded in response to RecA activation^58^. We aimed to quantify SOS induction using the pANO1::c*da*’ reporter plasmid, which expresses the fluorescent GFPmut3* protein from the SOS-inducible *cda* promoter^59^. MMC and GAT, a fluoroquinolone, were used as control compounds. GAT induces DNA damage and SOS as a secondary response to topoisomerase inhibition in bacteria, while it is not genotoxic to eukaryotic cells, which lack type II topoisomerases.

Green fluorescence was induced in response to GAT and MMC in growing cultures of UPEC CFT073 bearing pANO1::c*da*’. However, as the cultures approached the stationary phase, the drug-free control began to produce GFP as well. After 10 h, the fluorescence normalized to OD_600_ in the control culture surpassed that of the treated samples (Fig. S5A). Since we aimed to test SOS induction in non-growing cultures, it was essential to use a reporter with no leaky expression in the stationary phase. Therefore, we constructed the plasmid pAED2, based on the low copy-number plasmid pSC101, which contains the *cda*’ promoter in front of the mScarlet-I-encoding reporter gene. During a six-hour test, this reporter effectively lacked leaky expression and was induced by fluoroquinolones several thousand-fold, thus meeting our requirements (Fig. S5B).

We tested the SOS response to the anti-cancer agents in the medium where they showed higher activity (MMC, BLE, and ECH in 1:4 CA-MHB pH 7.4, PLI, SAT, and EVO in LPM pH5.5). MMC and SAT, as well as BLE, EVO, and ECH induced the SOS response to different extents when added to the growing cultures (Fig. 7). Induction of SOS by PLI was not detected (Fig. 7B). In accordance with their high MICs (Dataset 1), 20 μM PLI, EVO, and SAT cause only a slight growth inhibition compared to the effects of MMC, BLE and ECH (Fig S5 C,D). When added to stationary phase cultures, none of the compounds induced SOS (Fig. 7). This might indicate that the stationary phase bacteria are unable to induce a detectable SOS response because of the protein synthesis inhibition and are killed during the treatment. However, another possibility is that the bacteria are poisoned during the stationary phase treatment but are subject to DNA damage and initiate the SOS response only during regrowth after dilution into drug-free medium. We tested this possibility and observed no SOS induction despite the characteristic compound-specific regrowth delay (Fig. 7, Fig. S5E,F).

**Figure 7.**
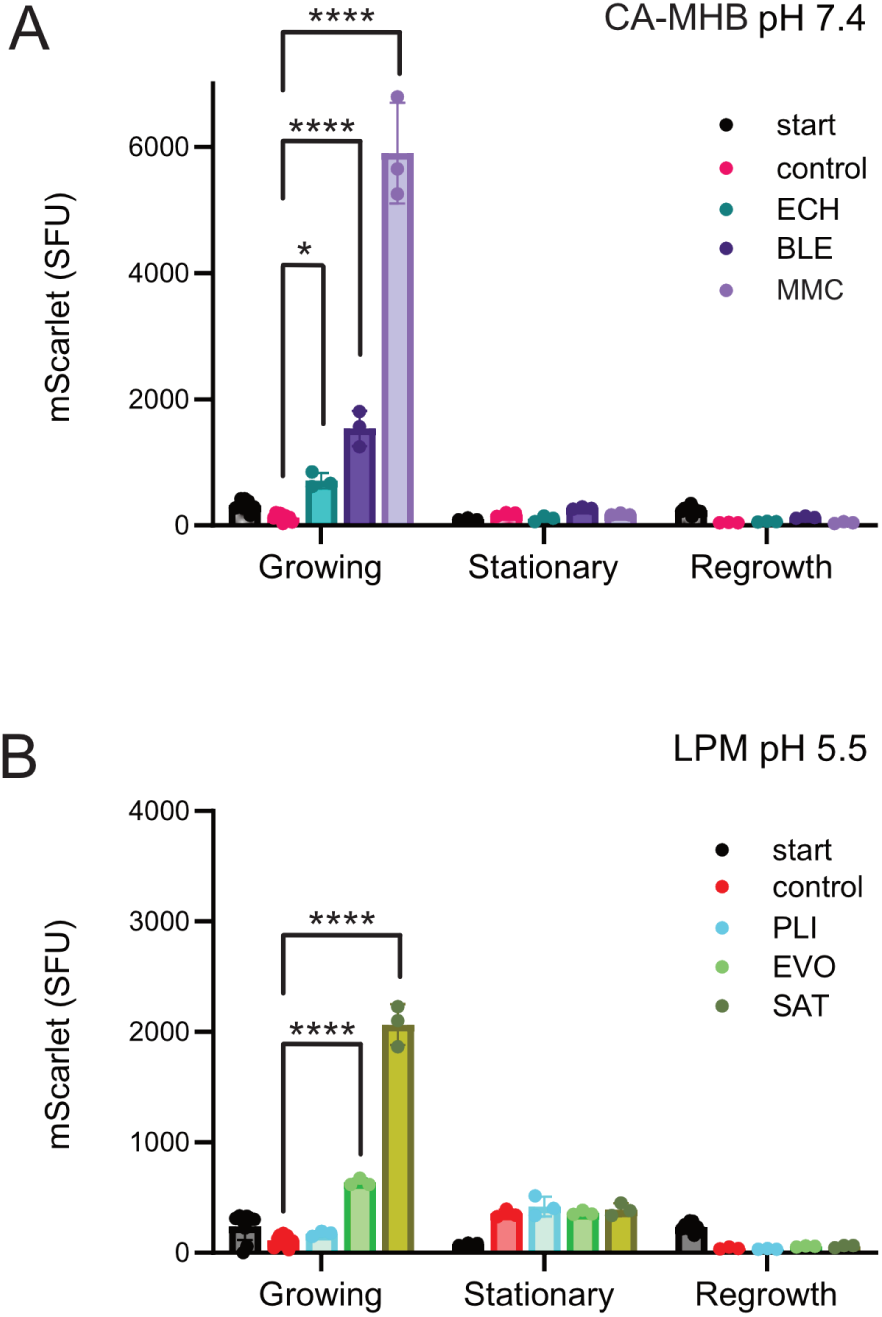
Induction of the SOS response by anti-cancer agents. UPEC CFT073 harboring pAED2 was cultivated in 1:4 CA-MHB (pH 7.4) to assess SOS response induction by echinomycin (ECH), bleomycin (BLE), and mitomycin C (MMC) (A), and in LPM (pH 5.5) to test plicamycin (PLI), evofosfamide (EVO), and satraplatin (SAT) (B). The expression of the mScarlet-I reporter, controlled by the SOS-inducible *cda*’ promoter, was evaluated by measuring the red fluorescence using a plate reader. Growing cultures were treated with 20 μM of each compound in a microplate, and fluorescence along with OD_600_ were measured. mScarlet fluorescence is shown at the start of incubation (start) and after 6 h for both the drug-free control (control) and treated samples. Stationary phase cultures were treated with 10 μM of each compound in test tubes for 24 h. Fluorescence and OD_600_ were measured at the start and end of incubation. After incubation, stationary phase bacteria were collected by centrifugation, washed, resuspended, and diluted 1:7 in CA-MHB. Regrowing samples were incubated in a microplate and fluorescence along with OD_600_ were measured. mScarlet fluorescence is shown at the start of the incubation (start) and when OD_600_ reached approximately 0.5 for both the drug-free control (control) and treated samples. Specific fluorescence units (SFU) were calculated by normalizing arbitrary fluorescence to cell density (AU/OD_600_). Values are presented as means ±SEM for n=3. Statistical significance was assessed using a one-way ANOVA followed by Dunnett’s test, comparing each treatment to the drug-free control. Four asterisks (****) denote Dunnett’s test p-values < 0.0001 and one asterisk (*) denotes p-value < 0.05.

### Evaluating hit compounds for growth inhibition of intracellular S. flexneri

Nonproliferating bacterial pathogens frequently inhabit intracellular environments, but establishing relevant infection models for high-throughput testing is challenging. Therefore, we firstly tested ability of the hit compounds to cross the eukaryotic cytoplasmic membrane barrier and inhibit growth of *Shigella flexneri* within the cytoplasm of the TC7 human colon adenocarcinoma cell line^60^. To this end, we developed a new, fluorescence-based method that enables real-time monitoring of growth of intracellular *Shigella*. We used *S. flexneri* 5a M90T strain producing the AfaE adhesin that enhances bacterial adhesion to intestinal epithelial cells and allows synchronization of invasion. The strain was further modified to constitutively express super-folder GFP (sfGFP) from the chromosomal Tn7 insertion site, which enables monitoring bacterial growth by fluorescence measurement. The epithelial cells were grown to confluence in 96-well plates and infected with bacteria at an MOI of 50. Following bacterial adhesion and invasion, the cells were incubated with varying concentrations of the hit compounds and 50 μg/ml gentamicin, which was added to inhibit extracellular bacteria but has no effect on intracellular bacteria. Intracellular bacterial growth was tracked by measuring green fluorescence every hour. At concentrations of antibacterials that permit bacterial growth, fluorescence intensity consistently increased over the 14-hour incubation period (Fig. 8A). To quantify the effect of the compounds, we calculated the IC_50_, representing the concentration at which intracellular bacterial growth was inhibited by 50%. This was determined from the endpoint fluorescence intensity (Dataset 3), with values ranging from baseline (no inhibition, fluorescence intensity similar to the drug-free control) to maximum effect (complete inhibition, with no increase in fluorescence) (Fig. 8B).

**Figure 8.**
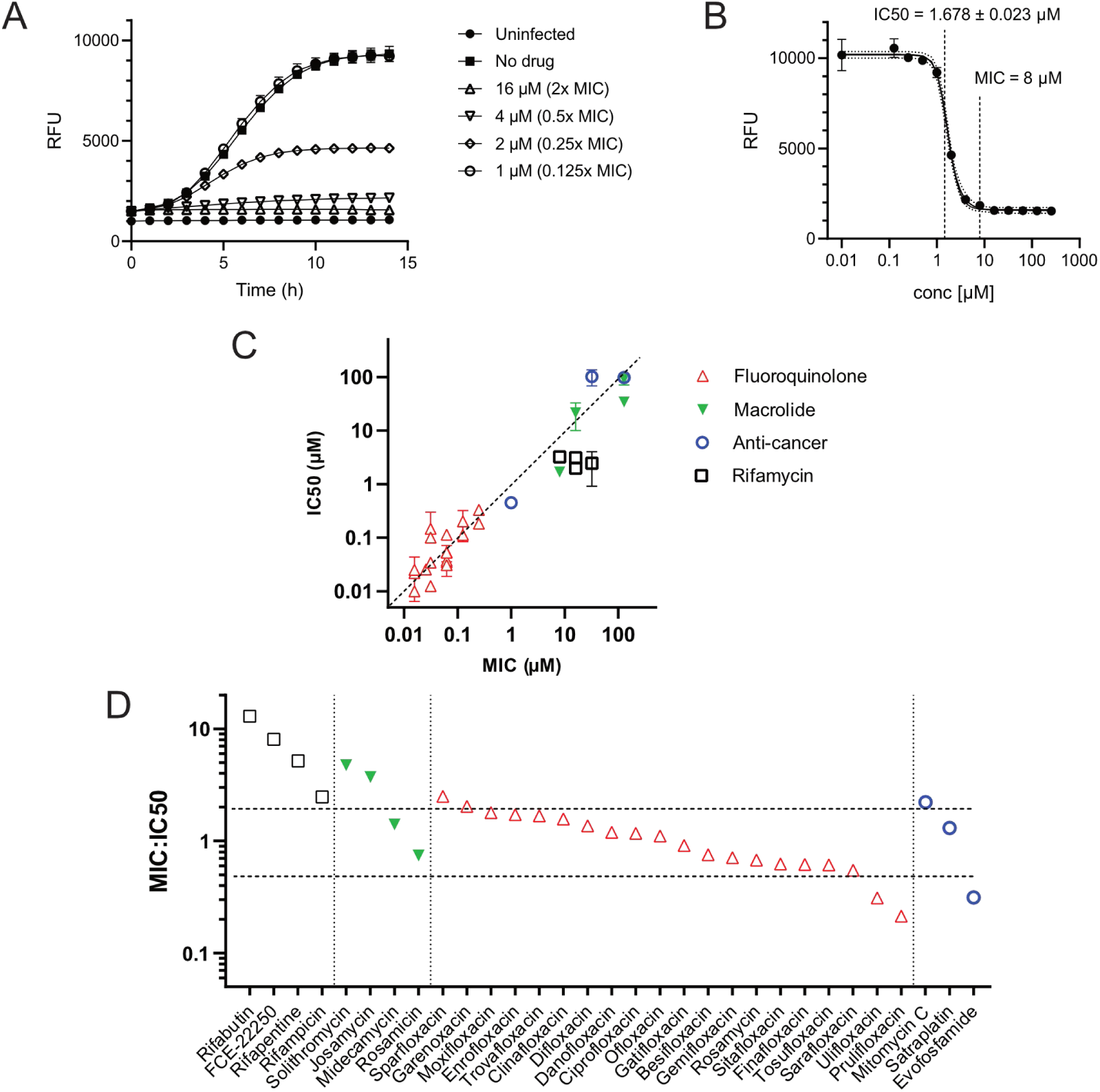
Inhibition of intracellular growth of *Shigella* in TC7 cells. TC7 cells were infected with *S. flexneri* constitutively expressing sfGFP and treated with various concentrations of the hit compounds. Gentamicin was used to eliminate extracellular bacteria and intracellular bacterial growth was monitored by measuring GFP fluorescence. RFU – relative fluorescence units. The symbols for antibiotic classes are shown next to panel C. **A.** Intracellular growth curves of *S. flexneri* in the presence of solithromycin at different concentrations as a representative example of a set of growth inhibition curves. Data are presented as means ± SD for n=3. **B.** Solithromycin inhibition curve. Intracellular growth inhibition was assessed by GFP fluorescence after 14 h treatment. Data are presented as mean ± SD for n=3. A five-parameter logistic equation was used for asymmetric sigmoidal curve fitting and IC_50_ determination in GraphPad Prism. For comparison, the MIC obtained by treating extracellular bacteria with the drug is also indicated. **C.** Comparison of MIC and intracellular IC_50_ values. Error bars represent 95% confidence intervals (CI). **D.** MIC:IC_50_ ratios with horizontal dashed lines indicating two-fold differences.

To evaluate the potentially toxic impact of the hit compounds on mammalian cell viability, we incubated TC7 cells with each drug in the absence of bacteria. At the MIC concentrations, satraplatin, chlorhexidine, alexidine, and valnemulin were toxic to epithelial cells (Fig. S6). Consequently, these compounds were excluded from further analysis. We observed that most compounds inhibited intracellular *Shigella*, with the exception of bleomycin, which was inactive (Dataset 1). A comparison of intracellular IC_50_ values with MICs revealed a strong correlation between intracellular and extracellular antibacterial activities (r = 0.79; 95% CI = 0.59 to 0.89; p < 0.0001). Fluoroquinolones with low MIC values were also among the most effective against intracellular *S. flexneri* (Fig. 8C). The MIC:IC_50_ ratios are generally close to one, with the exception of rifamycins, which showed notably higher ratios. All rifamycins have MIC values at least 2-fold higher than their IC_50_ values, with rifabutin exhibiting a an MIC:IC_50_ >10. Apart from rifamycins, solithromycin, josamycin, sparfloxacin, and mitomycin C also have an MIC:IC_50_ ratio >2, indicating enhanced activity against intracellular bacteria. In contrast, ulifloxacin, prulifloxacin, and evofosfamide display a slightly weaker intracellular activity, with MIC:IC_50_ ratios below 0.5 (Fig. 8D).

### The antibacterial effect of semapimod depends on conditions of cultivation and treatment

Recently, results from a similar screen conducted in conjunction with a deep learning-based virtual screen were reported by the Collins lab^61^. Semapimod, an anti-inflammatory drug with newly identified anti-membrane activity, was identified as a novel compound effective against stationary-phase *E. coli* and *Acinetobacter baumannii*^61^. Although semapimod was present in our compound library, it was not among our hits. We hypothesized that the discrepancy could be due to differences in key experimental setup. Zheng and colleagues^61^ grew stationary phase cultures of *E. coli in* 1% LB diluted in phosphate-buffered saline (PBS), similarly to a study where an overnight *S. aureus* MRSA culture was diluted 1:100 in PBS and then treated with antibiotics^62^. Additionally, they used a different *E. coli* strain (BW25113, a K12 derivative), a different screening concentration (50 μM vs 20 μM used here), as well as a 200-fold dilution into the drug-free growth medium, which was LB (we used a 2,500-fold dilution into CA-MHB pH7.4). The authors also observed that the activity of semapimod depends on the divalent cation concentration, as the addition of 21 mM Mg²⁺ abolished its activity. To investigate the causes underlying the apparent discrepancy in our findings, we tested semapimod against stationary phase cultures of both *E. coli* strains, combining different elements of these two protocols.

As shown in Fig. 9, the medium used for growing the stationary phase culture – which also served as the environment during treatment – is likely the primary factor impacting semapimod activity. Semapimod is strongly bactericidal (Fig. 9A) and delays regrowth (Fig. 9B, C) only when tested in 1% LB in PBS, but not in 1:4 CA-MHB. The other factors have relatively minor effects. Thus, the use of 1% LB in PBS by Zheng and colleagues^61^, compared to our use of 1:4 CA-MHB, explains the conflicting results of the screens.

**Figure 9.**
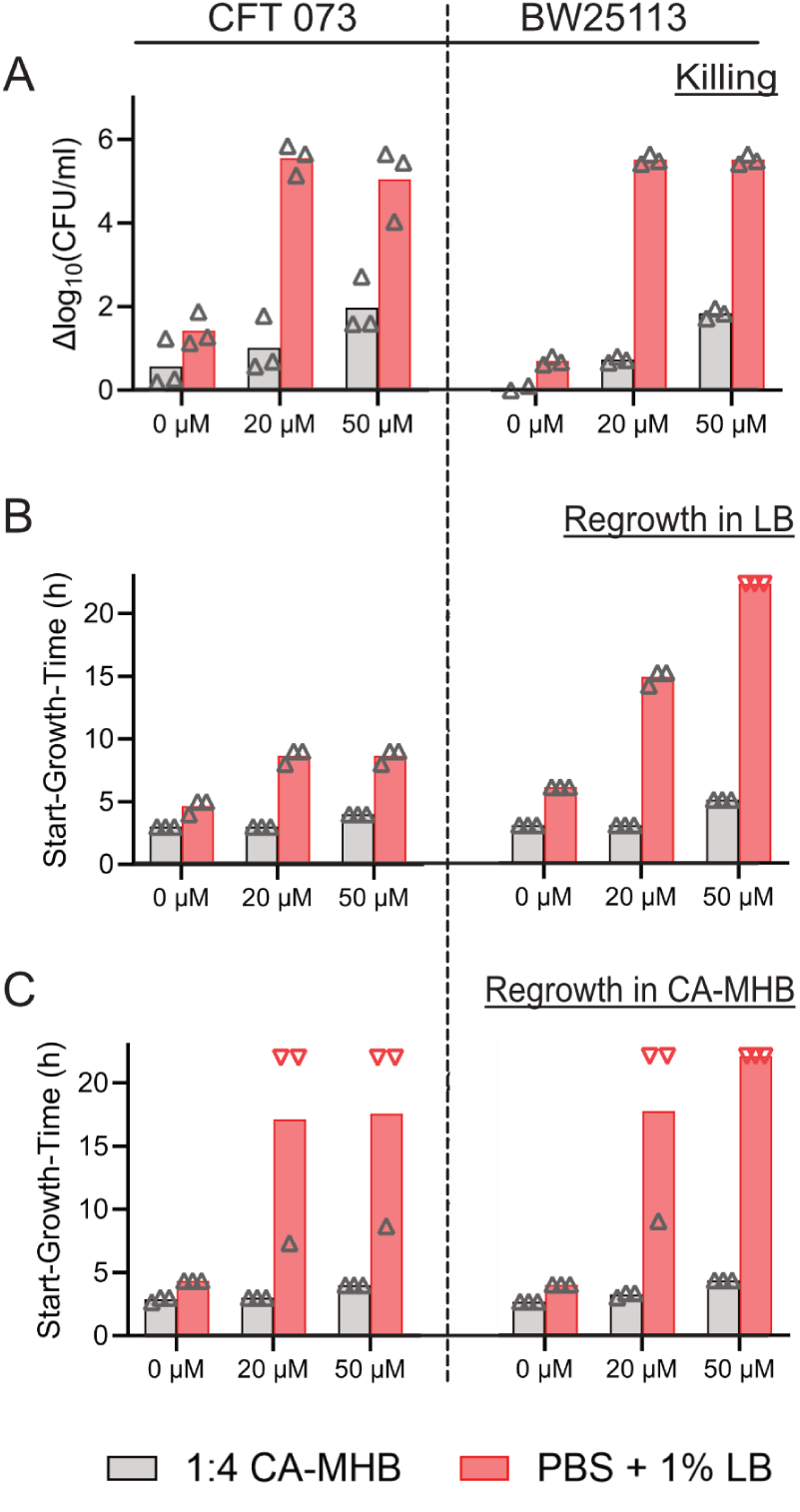
Bactericidal effect of semapimod on non-growing *E. coli* depends on a phosphate buffer-based culture medium. E. *coli* strains CFT073 and BW25113 were cultivated in either 1:4 CA-MHB pH 7.4 (gray columns) or PBS supplemented with 1% LB (pink columns) for 24 h, followed by treatment with 20 µM or 50 µM semapimod for an additional 24 h. **A.** To estimate survival, samples were serially diluted and spot-plated on LB agar. The differences in CFU/ml before and after treatment are presented. The average log bacterial densities before treatment (log_10_CFU/ml ± SEM) were 9.97 ± 0.63 for CFT073 grown in 1:4 CA-MHB, 9.49 ± 0.01 for BW25113 grown in 1:4 CA-MHB, 7.74 ± 0.05 for CFT073 grown in PBS + 1% LB, and 7.49 ± 0.07 for BW25113 grown in PBS + 1% LB. Red symbols represent individual experiments where bacterial counts fell below the limit of detection (no colonies from the undiluted sample). The maximum detectable decrease in CFU/ml for these samples is indicated and used for calculating average Δ log_10_CFU/ml. **B, C.** To assess regrowth ability, samples were diluted 200-fold into LB broth (B), and 2,500- fold into CA-MHB (C). OD_600_ was measured periodically for 22 h post-dilution. The time at which cultures reached an OD_600_ threshold of 0.12 is reported as the Start-Growth-Time (SGT). Red symbols represent individual experiments where cultures did not reach the threshold during 22 h. For these samples, SGT of 22 h is indicated and used for calculating average SGT. Data are presented as means for n=3.

## DISCUSSION

The development of antibiotics specifically targeting non-growing bacteria and chronic bacterial infections has been largely neglected. As a first step to address this gap, we conducted a drug-repurposing screen to identify antimicrobial agents that are primarily effective against quiescent Gram-negative pathogens. Importantly, we focused on compounds that have already undergone or are currently undergoing clinical trials, ensuring that their safety and pharmacokinetic properties are at least partially characterized. We also sought compounds active against Gram-negative pathogens residing in acidified intracellular compartments. This screen led to the identification of several antimicrobials with previously unrecognized potent activity against non-growing bacterial pathogens. While previous screening projects aimed at finding drugs against dormant bacteria have yielded somewhat different results – likely due to technical variations in screening and follow-up assays or differing criteria for prioritizing hit compounds^32,36,37,61^ – our approach offers new candidates that warrant testing against larger panels of clinical isolates.

A previous repurposing screen aimed at identifying bacterial growth inhibitors found that 24% of drugs targeting human cells, spanning all therapeutic classes, inhibited the growth of certain gut bacterial strains^63^. In contrast, we found that compounds effective against non-growing bacteria in this study were limited to a few antibiotic classes, with only select drugs within these classes demonstrating activity. For instance, among the quinolones included in the library, less than half showed bactericidal activity against non-growing bacteria. Among the 11 macrolides in the SPECS collection, five were active, while six—including widely used erythromycin, clarithromycin, and azithromycin – were not. Among the 14 DNA-alkylating agents, only three – mitomycin C, satraplatin and evofosfamide—exhibited activity against non-growing UPEC, and out of the 19 annotated DNA synthesis inhibitors, only bleomycin demonstrated such activity. As compounds within the same class target the same biological process, the formation of these two subgroups cannot be attributed to differences in target activity or corruption. Instead, this variation is likely due to different permeability or uptake of these drugs into non-growing and growing bacteria.

Additionally, we identified several compounds with high MIC values, indicating poor efficacy against growing bacteria, which were nonetheless highly effective against the nongrowing cells of the same strain. For instance, 0.5 µM solithromycin, a macrolide antibiotic of the ketolide sub-class, killed about four logs of stationary phase *P. aeruginosa*, despite its MIC being 80 μM. The discrepancy, once again, is likely attributable to differences in drug uptake between growing and non-growing bacterial cells. In contrast, telithromycin, another ketolide included in the library, showed no activity. The enhanced uptake of solithromycin into non-growing bacteria may be related to the presence of a fluoride atom, which is absent in telithromycin. Based solely on its MIC, solithromycin would not be considered a viable drug candidate against *P. aeruginosa*, underscoring the need for specialized testing procedures to identify antibiotics effective against persistent infections.

The different bactericidal activity of solithromycin against *P. aeruginosa* and *E. coli* may be attributed to variations in their ribosomal structures and, potentially, the slower dissociation of the drug from the *P. aeruginosa* ribosome. While bacteriostatic and bactericidal macrolides bind to the ribosome with comparable affinity, bactericidal drugs exhibit significantly slower dissociation. This prolonged inhibition of translation can damage bacterial cells, ultimately leading to death^64^. The slow dissociation of ketolides is attributed to their extended alkyl-aryl side chain, which establishes additional interactions with the ribosome. This side chain acts as a potential pharmacophore, offering opportunities to enhance the bactericidal properties of the drug.

In conclusion, purposefully modifying drugs to improve their penetration into non-growing bacteria and enhance their bactericidal effect could be a promising approach for developing agents against persistent infections. This strategy may enable the development of drugs specifically targeting non-growing bacteria (antiNG-antibiotics) and allow their combination with antibiotics that target actively growing pathogens (antiG-antibiotics, such as beta-lactams) during therapy. Notably, effective antiNG-antibiotics should possess high MIC values to avoid inhibiting bacterial proliferation and thereby preserving the bactericidal activity of the antiG- antibiotic.

A significant challenge in developing drugs for dormant pathogens is managing toxicity^36,61,65^. Half of the hits of our screen were fluoroquinolones, which have drawn increasing attention due to their severe adverse effects, and are no longer recommended as first-line antibiotics^66^. Sitafloxacin and clinafloxacin, which were among the most effective against non-growing bacteria, carry risks due to their extra halogen atoms, which enhance the penetration of lipid membranes but also increase undesirable accumulation in adipose tissue. Oral sitafloxacin is currently approved in Japan and Thailand, while clinafloxacin was withdrawn from development due to phototoxicity, a common issue among fluoroquinolones with a halogen at the C-8 position^67^. Topical application can mitigate the systemic toxicity of fluoroquinolones, as demonstrated by gatifloxacin, which is safely used for eye infections despite its oral form being banned^68^. To assess an antibiotic’s suitability for the topical treatment of chronic infections, such as chronic wound infections, targeted drug development and specialized testing are necessary.

We successfully employed our new GFP-based method to monitor the intracellular growth of *Shigella* upon challenge with drugs. All hit compounds, except those that were toxic to TC-7 cells or that could not be tested because of their high extracellular MIC values, were evaluated for their ability to inhibit intracellular *Shigella*. All but one compound (bleomycin) demonstrated activity, indicating they are able to permeate the cytoplasm of epithelial cells. However, it remains unclear whether rifapentine, solithromycin, and other drugs with higher MIC-to-intracellular IC_50_ ratios accumulate in the cytoplasm of TC-7 cells, or if intracellularly located *S*. *flexneri* is more sensitive to these drugs compared to its sensitivity in TSB broth.

Notably, our screening did not identify any non-antibacterial human drugs with generally new mechanisms of antibacterial activity. The findings included certain anti-cancer compounds with known nonspecific DNA-modifying activity, as well as the DNA-intercalating agents echinomycin and plicamycin, which have the potential to interfere with both transcription and replication. Although these compounds may have preferential targets in tumor cells, they have broad mechanisms of action on other cell types, which can account for their antibacterial properties. The targets of these compounds are common to both eukaryotic/cancer cells and bacteria. For instance, MMC is known for its anti-persister activity^69^, and, when used in combination therapy, is effective *in vivo* against multidrug-resistant (MDR) *P. aeruginosa* ^70^. The antimicrobial activity of DGS was identified through a repurposing screen^71^ and it is effective against multidrug-resistant *S. aureus* by modifying several essential cysteines and altering multiple pathways^47^. Echinomycin is active against *S. aureus*, including methicillin-resistant strains (MRSA)^72^. Testing the extent and timing of bacterial DNA damage induced by anti-cancer compounds did not provide a clear understanding of their activity against nongrowing bacteria. However, it appears that the echinomycin and plicamycin may be less genotoxic than the other DNA-targeting agents and warrant further investigation against persistent infections.

In conclusion, applying the dilution-regrowth screen enabled us to identify several promising compounds with potential activity against persistent Gram-negative infections, meriting continued study.

## METHODS

### Bacterial strains and plasmids

*Escherichia coli* CFT073, *Pseudomonas aeruginosa* DSM1117 (also known as Boston 41501, ATCC 27853), and *Staphylococcus aureus* DSM2569 were obtained from DSMZ (Deutsche Sammlung von Mikroorganismen und Zellkulturen GmbH, Braunschweig, Germany). *E. coli* BW25113 was from Coli Genetic Stock Center (CGSC). pANO1::c*da*’^59^ was a gift from Dr. Lars Hestbjerg Hansen. pAED2, a low-copy, ampicillin-resistant (AmpR) plasmid based on the pSC101 backbone, was constructed using circular polymerase extension cloning (CPEC)^73^ in two steps. First, the *trpT* terminator region from pSC101-CAM-bioreporter^74^ was inserted between *gfp*-mut2 and *mScarlet*-I genes in pSC101-GFPmut2-mScarlet-I^75^, resulting in plasmid pAED1. Next, the *cda* promoter from pANO1::*cda*’ was inserted upstream of the mScarlet-I gene, resulting in pAED2. Due to difficulties amplifying the *ampR* gene, the vector was amplified in two fragments. The primers used for cloning are listed in Table S1. *Shigella flexneri* 5a M90T contained a chromosomal insertion of constitutively expressed super-folder *gfp* gene at the attn∷Tn7 site, downstream the *glmS* gene, and was constructed as detailed previously^76,77^. *Shigella flexneri* M90T expressed the adhesin AFA-I (also known as AfaE^78,79^) from a plasmid conferring spectinomycin resistance.

### Dilution-regrowth screen

The high-throughput screening (HTS) procedure was based on the protocol by Hazan and colleagues^15^ and conducted at the Chemical Biology Consortium Sweden (CBCS) screening facility. Compounds from the Prestwick Chemical Library (1,200 compounds) and the Specs Repurposing Library (5,254 compounds) were dispensed using the Echo® (Labcyte) system directly into 384-well plates; 200 nl of 10 mM solution in DMSO was added per well. Each plate included eight wells with 200 nl of 10 mM gatifloxacin, finafloxacin, and kanamycin solutions, along with eight wells containing 200 nl of DMSO as controls. An overnight culture (∼16 h) of *E. coli* CFT073 was grown in LB broth, then diluted 1:100 into either 1:4 diluted CA-MHB medium buffered with 40mM HEPES (pH 7.4) or into LPM medium ^25^ supplemented with 49 µM MgCl_2_ and buffered with 30 mM MES (pH 5.5). The cultures were incubated for 24 h in non-baffled Erlenmeyer flasks at 37°C with shaking at 200 rpm. After incubation, 100 μl of the culture was dispensed into each well of the 384-well plates for screening at a final concentration of 20 μM. The plates were sealed with parafilm and incubated at 37°C for 24 h in closed humidified boxes, with wet paper towels added to prevent evaporation. After treatment, samples were first diluted 50-fold, using Beckman NX^P^ robot to transfer 2 μl of the sample to 98 μl of phosphate buffered saline (PBS), followed by a second 50-fold dilution into CA-MHB using the same volumes, resulting in a total 2500-fold dilution. The plates were incubated at 37°C for 6 h in humidified boxes, then OD_600_ was measured using a microplate reader. Compounds were considered active if the OD_600_ < 0.1. Initially, six plates were normalized using the drug-free control, but since this normalization had no effect on the results, it was later omitted.

For hit validation, bacteria were cultivated, and the dilution-regrowth assay was conducted following the same protocol used in the primary screen. Hit compounds were dispensed into 384-well plates using the Echo® (Labcyte) system, with 200, 100, or 50 nl of a 10 mM solution in DMSO added per well, yielding final screening concentrations of 20, 10, and 5 μM, respectively. OD_600_ values of regrowing samples were normalized to the mean OD_600_ of the drug-free control. The testing was conducted in two replicates. Compounds were considered active if their OD_600_ was at least 50% lower than that of the drug-free control at one or more concentrations in both replicates.

For screening assay validation in pilot experiments, bacteria were cultivated as later in the screening assays. Aliquots of 180 μl from the stationary-phase culture were transferred to 96-well plates, and 20 μl of a 200 μM antibiotic solution (gatifloxacin or finafloxacin) in 10% DMSO or 10% DMSO alone for drug-free controls were added. After 24 h treatment, 4 μl of the sample was diluted in 196 μl of PBS, followed by a second dilution of 4 μl into 196 μl of CA-MHB. For assay validation in 384-well plates, 90 μl of the stationary-phase culture was mixed with 10 μl of the 200 μM antibiotic solution or 10% DMSO. After the treatment, 2 μl of the sample was diluted in 98 μl of PBS, followed by a second dilution of 2 μl into 98 μl of CA- MHB. To monitor regrowth, the plates were incubated in a microplate reader (BioTek Synergy Mx) at 37°C with continuous shaking, and OD_600_ was recorded every 20 minutes. For regrowth validation in standing plates, the plates were incubated at 37°C in humidified boxes, and OD_600_ was measured after 5, 6, 7, and 8 h. The assays were carried out in triplicates.

### Dose-response analysis using the dilution-regrowth assay

For UPEC, dose-response analysis was conducted at the CBCS screening facility. Bacteria were cultivated, and the dilution-regrowth assay was performed according to the HTS protocol. Compounds were dispensed into 384-well plates using the Echo® (Labcyte) system, and regrowth was assessed by measuring OD_600_ after 6 h of incubation at 37°C in humidified boxes. Three biological replicates were performed.

For *P. aeruginosa* and *S. aureus*, dose-response analysis was conducted in 96-well plates with a 200 μl assay volume, following the procedure used for screening assay validation experiments. Aliquots of the 39 compounds active against non-growing UPEC were obtained from SelleckChem and MedChem Express. Stock solutions were prepared and stored according to the manufacturers’ instructions, then diluted in 10% DMSO before use. Cultures were incubated for 24 h in 1:4 diluted CA-MHB medium under the same conditions used for UPEC cultivation. In preliminary experiments, each compound was tested at a concentration of 20 μM in the dilution-regrowth assay with 4 technical replicates. Regrowth was monitored using a microplate reader at 37°C with continuous shaking, and OD_600_ was recorded every 20 min. The time at which OD_600_ exceeded the threshold of 0.12 was recorded as the Start_Growth-Time. Based on these results, 24 of the most active compounds were selected for further dose-response analysis, which was performed in three biological replicates using the same procedure.

### Dose-dependent killing analysis

For survival analysis, bacteria were cultivated as described for the dose-response dilution-regrowth assay. Assays with UPEC were conducted at the CBCS screening facility using 96- well plates with a 200 μl sample volume, and compounds were dispensed using the Echo® (Labcyte) system. *P. aeruginosa* and *S. aureus* samples were treated according to the protocol used for the dilution-regrowth experiments. Treated samples were serially diluted 10-fold in PBS, and 10 μl drops were spot-plated onto rectangular LB agar plates using either a Beckman NXP robot (for UPEC) or an Integra VIAFLO96 electronic pipette (for *P. aeruginosa* and *S. aureus*). The LB agar plates were incubated overnight at 37°C and scanned. Colonies were counted at the spots of the lowest dilution where individual colonies could be accurately distinguished. Each colony was manually marked on a separate image layer using the GNU Image Manipulation Program, and counted using NIST’s Integrated Colony Enumerator (NICE 1.4) software ^80^. The experiments were performed in three biological replicates.

### Susceptibility testing

Minimal Inhibitory Concentrations (MICs) were determined using a standard microdilution assay protocol ^80^. CA-MHB medium buffered with 40 mM HEPES (pH 7.4) was used to determine the MICs for UPEC, *P. aeruginosa*, and *S. aureus*. For compounds with high MIC values, the measurements were repeated in 1:4 diluted CA-MHB (40 mM HEPES, pH 7.4) and in CA-MHB (40 mM HEPES, pH 7.4) supplemented with 10 μg/ml phenylalanine-arginine β- naphthylamide (PaβN). MICs for UPEC were also determined in LPM buffered with 30 mM MES (pH 5.5). For *Shigella flexneri*, MICs were determined in tryptic soy broth (TSB). The assay was performed in triplicates.

### SOS response induction analysis

Overnight cultures of CFT073 harboring a reporter plasmid were grown in 3 ml of LB supplemented with 100 µg/ml carbenicillin (Cb), then diluted 1:100 into the medium used for testing. For validation of the pANO1::cda’ reporter, an overnight culture was diluted into CA- MHB (supplemented with 40 mM HEPES, pH 7.4, and 100 µg/ml Cb) and grown to an OD_600_ ≈ 0.5 at 37°C with shaking at 200 rpm in an Erlenmeyer flask. Aliquots of 200 μl were dispensed into a 96-well plate and treated with 0.1 and 0.5 μM gatifloxacin, mitomycin C, or left drug-free. Empty wells were filled with water. The plate was incubated in a microplate reader (BioTek Synergy H1) at 37°C with continuous shaking for 20 h. OD_600_ and GFPmut3* fluorescence (λ_ex_ = 485 ± 9 nm, λ_em_ = 508 ± 9 nm) were recorded every 10 minutes.

For pAED2 validation, the overnight culture was diluted into 3 ml of LB supplemented with 100 µg/ml Cb and incubated at 37°C with shaking at 200 rpm for 1h. Aliquots of 200 μl were dispensed into a 96-well plate and incubated for 6h with 0.1 and 0.3 μM gatifloxacin, ofloxacin, or left drug-free. OD_600_, GFPmut2 fluorescence (λ_ex_ = 485 ± 9 nm, λ_em_ = 508 ± 9 nm), and mScarlet fluorescence (λ_ex_ = 569 ± 9 nm, λ_em_ = 594 ± 9 nm) were measured using a BioTek Synergy H1 microplate reader. The experiment was performed in triplicate.

For analysis of the SOS induction by anti-cancer compounds, the overnight cultures were diluted either into 1:4 diluted CA-MHB (40 mM HEPES, pH 7.4) or LPM (30 mM MES, pH 5.5) that were supplemented with 100 µg/ml Cb. For analysis of the effect on growing bacteria, the cultures were incubated at 37°C with shaking at 200 rpm in Erlenmeyer flasks for 90 min, to an OD_600_ ≈ 0.5. Then, 180 μl aliquots were transferred to 96-well plates, and 20 μl of the 200 μM drug solution was added to achieve a final concentration of 20 μM. The plates were incubated in a microplate reader (BioTek Synergy H1) at 37°C with continuous shaking for 6 h. OD_600_ and mScarlet fluorescence were recorded throughout the incubation.

For analysis of the effect of anti-cancer compounds on stationary phase- and regrowing bacteria, the cultures were incubated for 24 h at 37°C with shaking at 200 rpm in Erlenmeyer flasks. For the drug treatment, 475 μl of the culture was mixed with 25 μl of a 200 μM drug solution to achieve a final concentration of 10 μM, which had been demonstrated to be effective against stationary-phase bacteria. The samples were incubated for 24 h at 37°C in standing 1.5 ml tubes. Before and after the drug treatment, 25 μl aliquots were taken and diluted by mixing with 175 μl PBS. OD_600_ and mScarlet fluorescence were recorded using a microplate reader (BioTek Synergy H1).

After the treatment, stationary-phase bacteria were harvested by centrifugation at 5000×g for 5 min at 20°C, washed twice with PBS, and resuspended in 475 μl PBS. Then, 25 μl aliquots were transferred to 96-well plates, and 175 μl CA-MHB was added. The plates were incubated in a microplate reader at 37°C with continuous shaking for 14 h. OD600 and mScarlet fluorescence were recorded throughout the incubation. Specific fluorescence was calculated by dividing fluorescence intensity (in relative fluorescence units, RFU) by optical density. The experiments were performed in three biological replicates.

### Cultivation of the TC7 intestinal epithelial cells

TC7 cells (a human cell line derived from Caco-2 colon adenocarcinoma cells ^60^) were grown in Dulbecco’s Modified Eagle Medium (DMEM; Gibco, catalog number 21885-025) supplemented with 10% heat-inactivated serum, 1x amino acid solution prepared from Gibco Minimum Essential Medium Non-Essential Amino Acids (100x) (catalog number 11140-035), and 100 U/ml penicillin-streptomycin (Gibco, catalog number 15140-122). The cells were split using trypsin, seeded in 96-well plates for infection, and grown to confluence, corresponding to approximately 40,000 cells per well.

### In vitro *infection of cell culture and susceptibility testing of intracellular* S. flexneri

A single colony of *S. flexneri* was inoculated into TSB supplemented with 50 μg/ml spectinomycin and incubated overnight at 37°C with agitation. The overnight culture was diluted 1:200 in fresh TSB, and the bacteria were grown at 37°C until the OD_600_ reached 0.3– 0.4, corresponding to approximately 2 × 10⁷ CFU/ml. Bacteria were pelleted by centrifugation, washed twice with sterile PBS, and resuspended in infection solution (serum-free DMEM supplemented with 20 mM HEPES, pH 7.4). Just prior to infection, the medium was withdrawn, and the cells were washed twice with sterile PBS. The bacterial inoculum (MOI 50, 2 × 10⁷ CFU/ml) was added to all wells except the no-infection control. The plates were incubated at room temperature for 15 min to allow bacterial adhesion, followed by a 1 h incubation at 37°C with 5% CO_2_ to facilitate bacterial invasion.

Two-fold serial dilutions of the drugs were prepared in infection solution containing 50 μg/ml gentamicin in a 96-well plate. After the invasion period, the bacterial inoculum was removed, and the serial dilutions of the drugs were added to the wells, except for the no-drug control. The plates were incubated in a Spark multi-mode plate reader (Tecan) at 37°C with 5% CO_2_ for 12 h. Green fluorescence from intracellular bacteria was recorded hourly for 15 h at λ_ex_ = 475 nm and λ_em_ = 510 nm, measuring 12 distinct regions of each well. The experiments were performed in triplicates on separate days.

### Determination of cellular viability upon drug exposure

TC7 cells were grown to confluence in transparent 96-well cell culture plates (TPP) in the same cell culture medium used for the *in vitro* infection assay. Just prior to the experiment, the medium was discarded, and the cells were washed twice with sterile PBS to remove any traces of medium or dead cells. A total of 100 μl of infection solution containing MIC concentrations of individual drugs was added to duplicate wells. The plate was incubated at 37°C in a CO₂ incubator for 12 h, under the same conditions as the infection assay.

The 3-(4,5-dimethylthiazol-2-yl)-5-(3-carboxymethoxyphenyl)-2-(4-sulfophenyl)-2H- tetrazolium (MTS) assay was performed using the CellTiter 96® AQueous One Solution Cell Proliferation Assay kit (Promega), according to the manufacturer’s instructions. Briefly, 20 μl of CellTiter 96 AQueous One Solution Reagent was added to each well, and the plate was incubated for 2 h at 37°C in a CO₂ incubator. The OD_490_ of the samples was measured using a Spark multimode plate reader (Tecan). The raw values were normalized to the control wells incubated with infection buffer only. The experiment was performed in duplicate.

For compounds that strongly absorb light at 490 nm (FCE-22250, rifampicin, mitomycin C, rifabutin, rifapentine), a lactate dehydrogenase (LDH) release assay was performed instead of the MTS assay to assess cytotoxicity. The LDH assay was conducted using the CytoTox-ONE Homogeneous Membrane Integrity Assay kit (Promega), according to the manufacturer’s instructions. Briefly, cells in positive control wells were lysed with 10 μl of 10x lysis solution. Then, 50 μl of supernatant from each well was transferred to a black 96-well microplate (Greiner), and 50 μl of CytoTox 96 Reagent was added to each sample. The plate was incubated in the dark for 10 min at room temperature, followed by the addition of 50 μl of Stop Solution. Fluorescence was measured at λ_ex_ = 560 nm and λ_em_ = 590 nm using a Spark multimode plate reader (Tecan). The raw values of the samples were normalized to the average of the total lysis controls, which was set to 100% cell death.

### Time-dependent effects of drug treatment

Bacteria were cultivated in 1:4 diluted CA-MHB (40 mM HEPES, pH 7.4) for 24 h. Aliquots were dispensed into a 96-well plate, and compounds were added at a 10 µM concentration. Samples were collected at designated time points, diluted 2500-fold as described for the dilution-regrowth assay, and incubated in standing plates within a humid box at 37°C. The OD_600_ of the samples was measured to assess regrowth using a microplate reader (BioTek Synergy H1) after 6 h for UPEC and 8 h for P. aeruginosa and S. aureus. The experiment was performed in triplicates.

### Semapimod assay

*E. coli* CFT073 and BW25113 were grown in 3 ml of LB at 37°C, shaking at 200 rpm for 16 h. The overnight cultures were then diluted either 1:100 into 40 ml of 1:4 diluted CA-MHB (40 mM HEPES, pH 7.4) or 1:10,000 into 40 ml of 1% LB in PBS and grown in 125 mL Erlenmeyer flasks for 24 h at 37°C, shaking at 200 rpm. Following this, 1 ml aliquots of each culture were transferred to 1.5 ml test tubes, and semapimod was added at final concentrations of 20 μM and 50 μM. DMSO was added to drug-free controls. To quantify live bacteria before treatment, a sample of each culture was serially 10-fold diluted into PBS, and 10 μL drops were spot-plated onto LB agar. The closed test tubes were incubated for 24 h at 37°C without shaking.

After treatment, the samples were diluted 2,500-fold in two steps: a 50-fold dilution into PBS, followed by a 50-fold dilution into CA-MHB in a 96-well plate with a total volume of 200 μl per well. The 96-well plate was then incubated in a BioTek Synergy H1 plate reader at 37°C with continuous shaking at medium speed for 22 h with a lid. OD600 was recorded every 20 minutes. Additionally, the treated samples were diluted 1:200 into LB in another 96-well plate and incubated in a BioTek Synergy Mx plate reader at 37°C without shaking for 22 h, with OD_600_ recorded every hour.

To determine the number of live bacteria after the treatment, the bacteria were pelleted, washed twice with PBS, serially diluted, and spot-plated. Plates were incubated and colonies counted as described previously. The experiment was performed in triplicates.

### Data analysis

The statistical analysis of all data, as well as EC_50_ and IC_50_ determination, was conducted using GraphPad Prism 10.2.3. Statistical significance of the drug effects was assessed using a one-way ANOVA followed by Dunnett’s test, comparing each treatment to the drug-free control. Statistical significance was set at P ≤ 0.05. A five-parameter logistic equation was used for asymmetric sigmoidal curve fitting and for EC_50_ and IC_50_ determination. The number of biological replicates and the meaning of the error bars are provided in the figure legends.

## ACKNOWLEDGMENTS

We thank Jelena Kiprovskaja for excellent technical assistance.

This work was financially supported by the Knut and Alice Wallenberg Foundation (project grant 2020-0037 to V.H.), the Swedish Research Council (Vetenskapsrådet) grants (2021- 01146 to V.H.), Cancerfonden (20 0872 Pj to V.H.), Göran Gustafsson Foundation for Research in Natural Sciences and Medicine (the Göran Gustafsson Prize to V.H.), Institutional support from MIMS (Vetenskapsrådet 2021-06602 to A.P), Kempe Stiftelserna (SMK-1859 and JCK-2031.3 to A.P), Svenska Sällskapet för Medicinsk Forskning (SSMF) postdoctoral grant (PD20-0022 to A.S.), Estonian Research Council grant (PRG335 to T.T.), MIBEst H2020-WIDESPREAD-2018-2020/GA 857518 (to T.T. and V.H.).The chemical libraries were distributed from CBCS Compound Center. CBCS is a national infrastructure funded by the Swedish Research Council (dr.nr.2021-00179) and SciLifeLab.

## AUTHOR CONTRIBUTIONS

N.K., T.T., V.H. and A.P. conceived the project. N.K. and S.B.F. performed the screening and analyzed the data. N.B., S.B.F., and N. K. performed the follow-up characterization of antimicrobials. A.S. and A.P. conceived and performed the *Shigella* and toxicity testing. N. Ch. A. performed the SOS response testing. J.A. and M.H. constructed the SOS reporter plasmid.

