## Supplementary Table 1, Supplementary Figures S1...S6. for "Antibacterial Compounds Against Non-Growing and Intracellular Bacteria"

**Supplementary Table 1.** CPEC primers

| **Primer** | **Sequence** | **Template** |
|---|---|---|
| Vector | | |
| term_vector_fwd | CAATATGGTGAGCAAGGGCGAGG | pSC101-GFP-mScarlet-I[1] |
| promoter_vector_rev | CTGTCAGGTCATTTCCAAGCTTGTCGA | |
| TestprimerAmp_rev | GACACGGAAATGTTGAATAC | pAED1 (this study) |
| AmpCPEC_fwd | GGAAGAGTATGAGTATTCAACATTTCCGTGTC | |
| *trpT* terminator insert | | |
| term_cpec_insert_fwd | CAACCGCAGTGAGTGAGTCTG | pSC101-CAM-bioreporter[2] |
| term_cpec_insert_rev | CCTCGCCCTTGCTCACCATATTGTGGTCAGTCATTTCCAAGCTTGTCGACCTGC | |
| *cda* promoter insert | | |
| pcda_insert_fwd | TCGACAAGCTTGGAAATGACCTGACAGCGCTCTTCGGCTTCGGTCA | pANO1::*cda*'[3] |
| pcda_insert_rev | CCTCGCCCTTGCTCACCATATTGCACCTCCTTGACTTTTAAAACAATGCGTTAAAAACAACAAAC | |


1. Hinnu, M., Putrinš, M., Kogermann, K., Kaldalu, N. & Tenson, T. Fluorescent reporters give new insights into antibiotics-induced nonsense and frameshift mistranslation. *Sci. Rep.* **14**, (2024).

2. Preem, L. *et al.* Monitoring of Antimicrobial Drug Chloramphenicol Release from Electrospun Nano- and Microfiber Mats Using UV Imaging and Bacterial Bioreporters. *Pharm. 2019, Vol. 11, Page 487* **11**, 487 (2019).

3. Norman, A., Hansen, L. H. & Sørensen, S. J. Construction of a ColD cda promoter-based SOS-green fluorescent protein whole-cell biosensor with higher sensitivity toward genotoxic compounds than constructs based on recA, umuDC, or sulA promoters. *Appl. Environ. Microbiol.* **71**, 2338–2346 (2005).

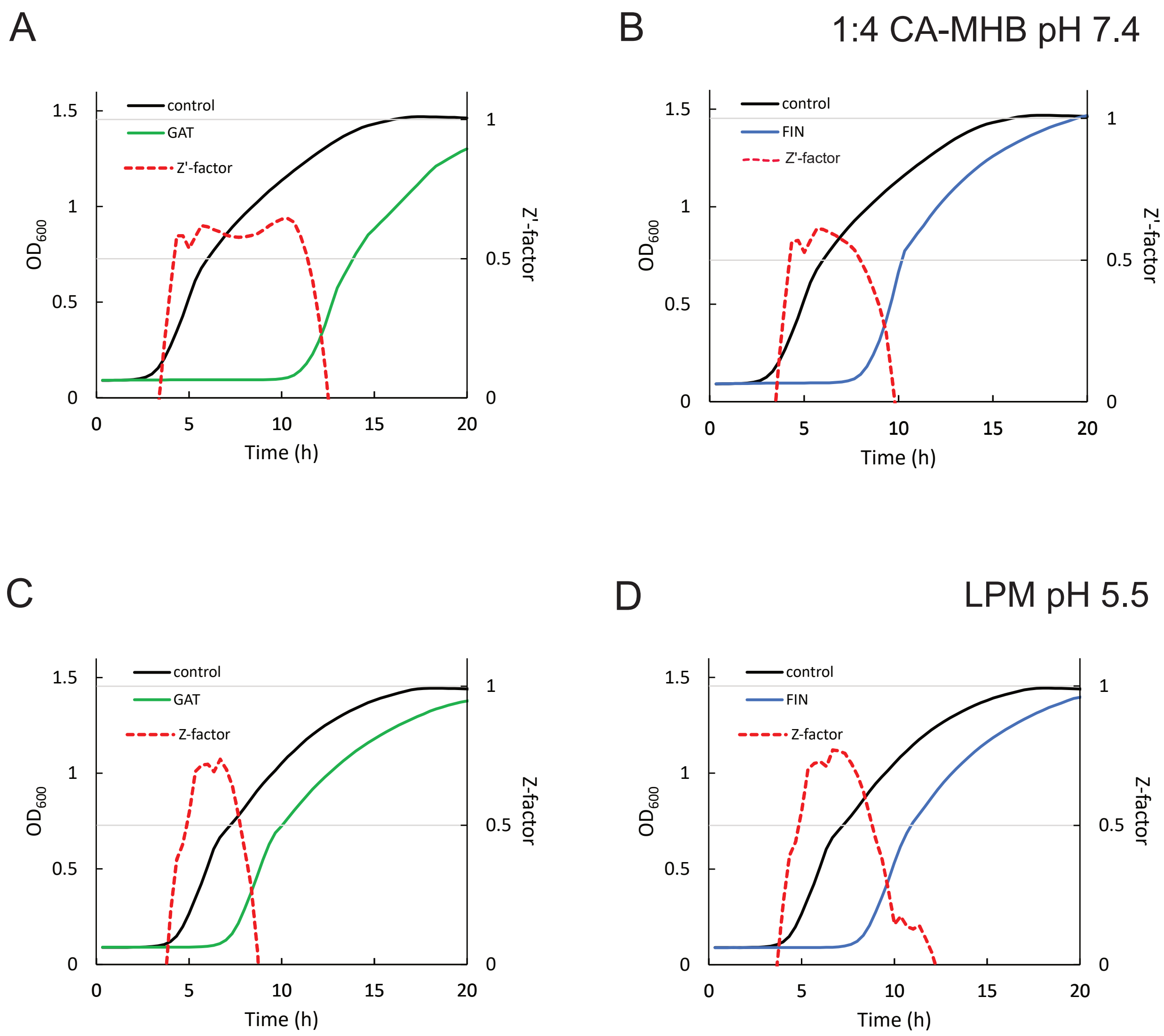


**Figure S1. Validation of the screening assay to identify compounds active against non-growing UPEC.**

UPEC CFT073 was cultivated in 1:4 diluted CA-MHB (pH 7.4) (A, B) or LPM (pH 5.5) (C, D) for 24 h. Cultures were then treated with 20 µM gatifloxacin (GAT) (A, C), finafloxacin (FIN) (B, D), or incubated without antibiotics as a control in a 96-well plate for 24 h. Regrowth was monitored by measuring $OD_{600}$ after a 2,500-fold dilution into CA-MHB. Each experiment included 12 technical replicates, with mean $OD_{600}$ values indicated by lines. OD600 readings were used to calculate the Z'-factor for the assay.

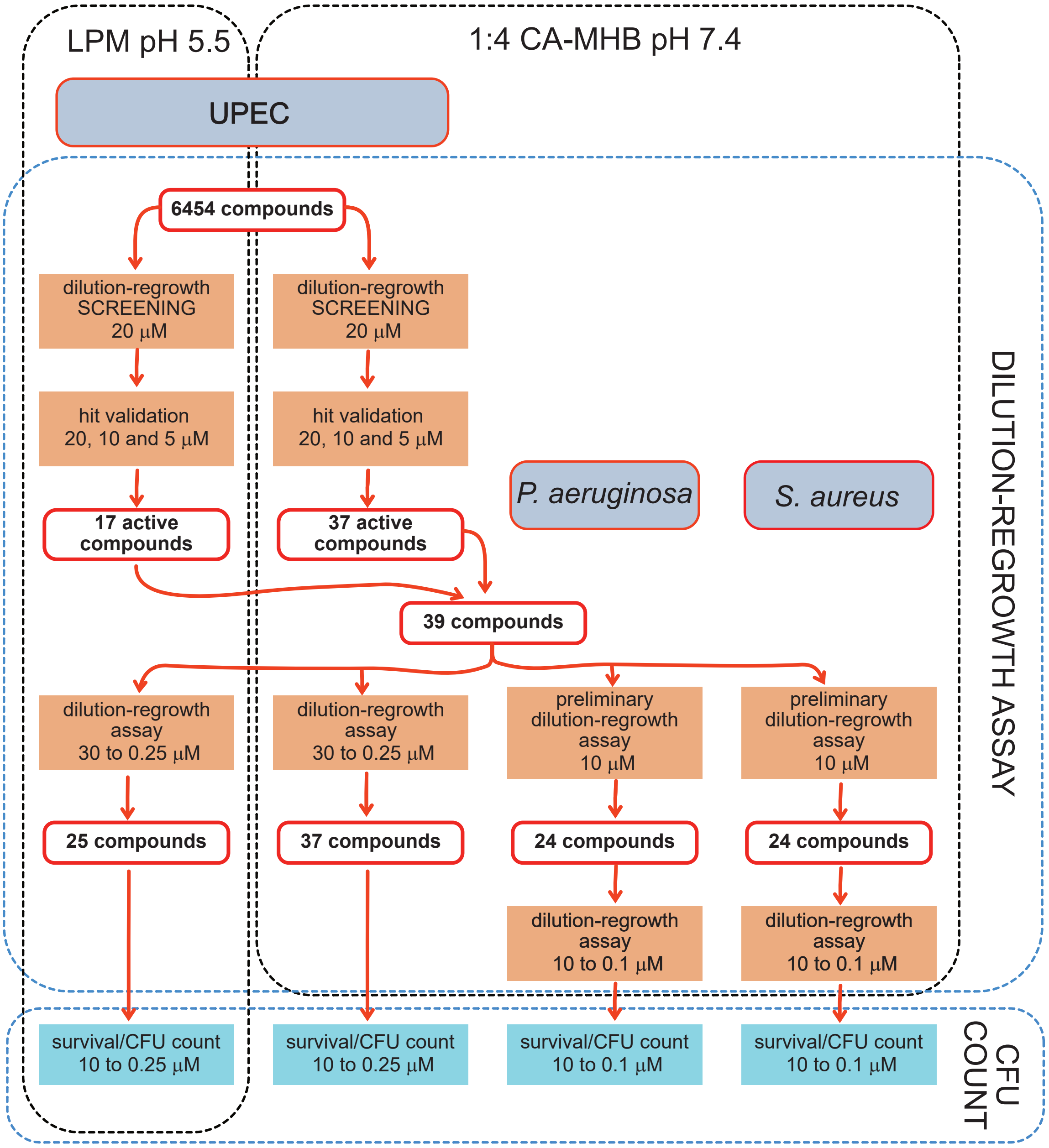


**Figure S2. Workflow for the screening and follow-up characterization of compounds active against non-growing bacteria.**

The screening of combined Prestwick and SPECS collections identified compounds that delayed the regrowth of stationary phase UPEC CFT073 after treatment in 1:4 diluted CA-MHB (pH 7.4) and LPM (pH 5.5). These hits were validated at three different concentrations. A total of 39 active compounds were tested at concentrations ranging from 30 to 0.25 µM for their ability to postpone UPEC regrowth in both 1:4 CA-MHB (pH 7.4) and LPM (pH 5.5). The verified active compounds were further tested for their bactericidal activity against UPEC in both media. The same set of 39 hit compounds was also tested against non-growing *Pseudomonas aeruginosa* DSM1117 and *Staphylococcus aureus* DSM2569. Based on preliminary dilution-regrowth assay results, the 24 most active compounds against each organism were identified and characterized for concentration-dependent regrowth delay and killing of non-growing *P. aeruginosa* and *S. aureus.*

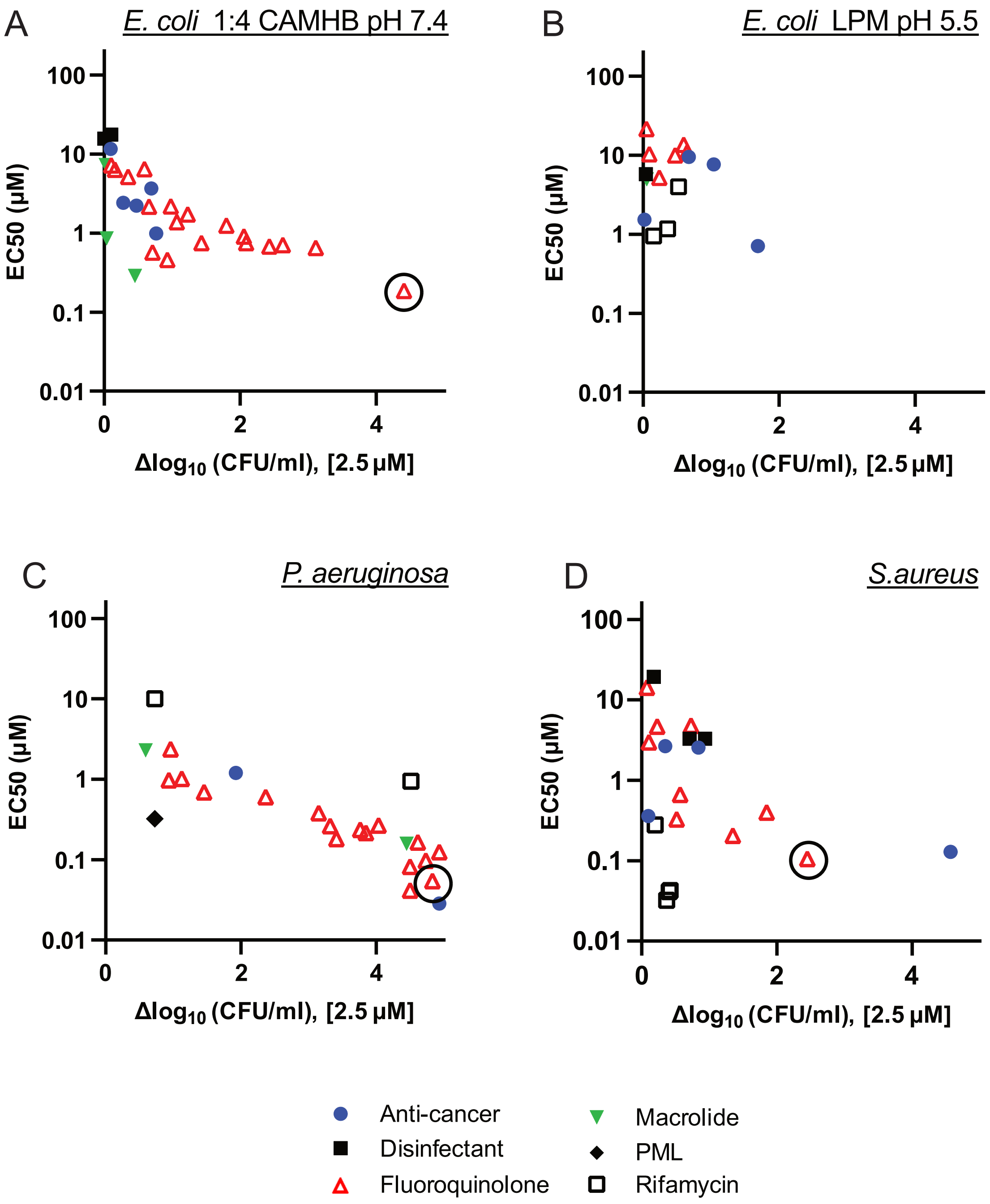


**Figure S3. Regrowth inhibition and bactericidal effects of hit compounds on non-growing bacteria.**

Bacterial cultures were grown for 24 h, followed by an additional 24 h treatment in 1:4 CA-MHB (pH 7.4) (A, C, D) or LPM (pH 5.5) (B). Regrowth was assessed by measuring $OD_{600}$ after a 2,500-fold dilution into CA-MHB. Regrowth inhibition is represented by EC50 values based on OD600 readings taken 6 h (A, B) or 8 hours (C, D) after dilution. Bacterial killing was evaluated by counting CFUs from samples treated with 2.5 µM of the compounds. The Δlog10 (CFU/ml) indicates the difference between drug-treated samples and the drug-free control. Sitafloxacin is highlighted with a circle.

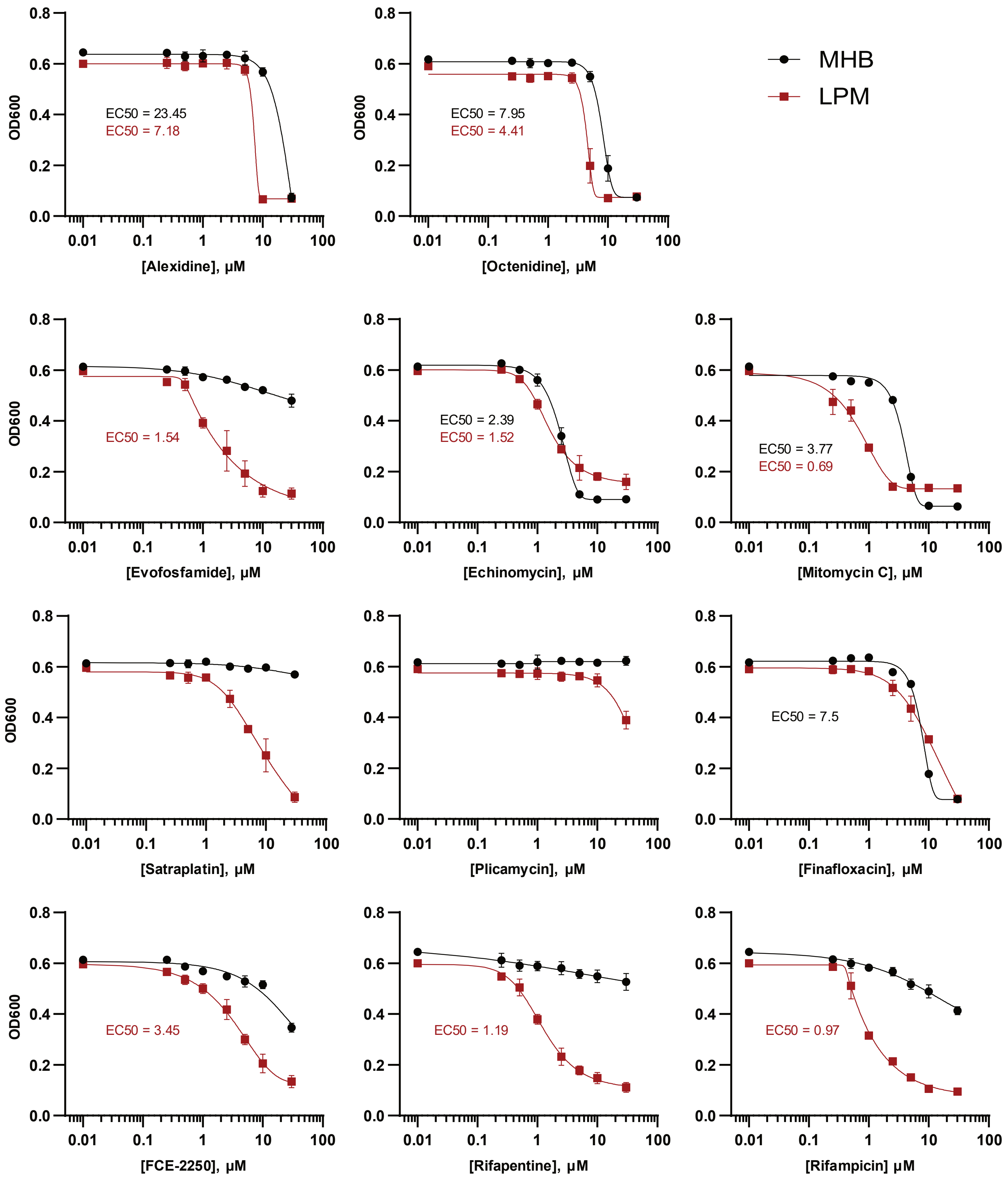


**Figure S4. Hit compounds with enhanced activity against non-growing UPEC in acidic medium**

Regrowth inhibition curves of the hit compounds with superior activity against non-growing UPEC in LPM (pH 5.5) (red) compared to 1:4 CA-MHB (pH 7.4) (black). Post-treatment regrowth inhibition was assessed by measuring $OD_{600}$ 6 h after a 2,500-fold dilution into CA-MHB. Data are presented as means ±SEM for n=3. A five-parameter logistic equation was used for asymmetric sigmoidal curve fitting and $EC_{50}$ calculation in GraphPad Prism. $EC_{50}$ calculations were omitted when the $OD_{600}$ values did not form a complete sigmoidal curve.

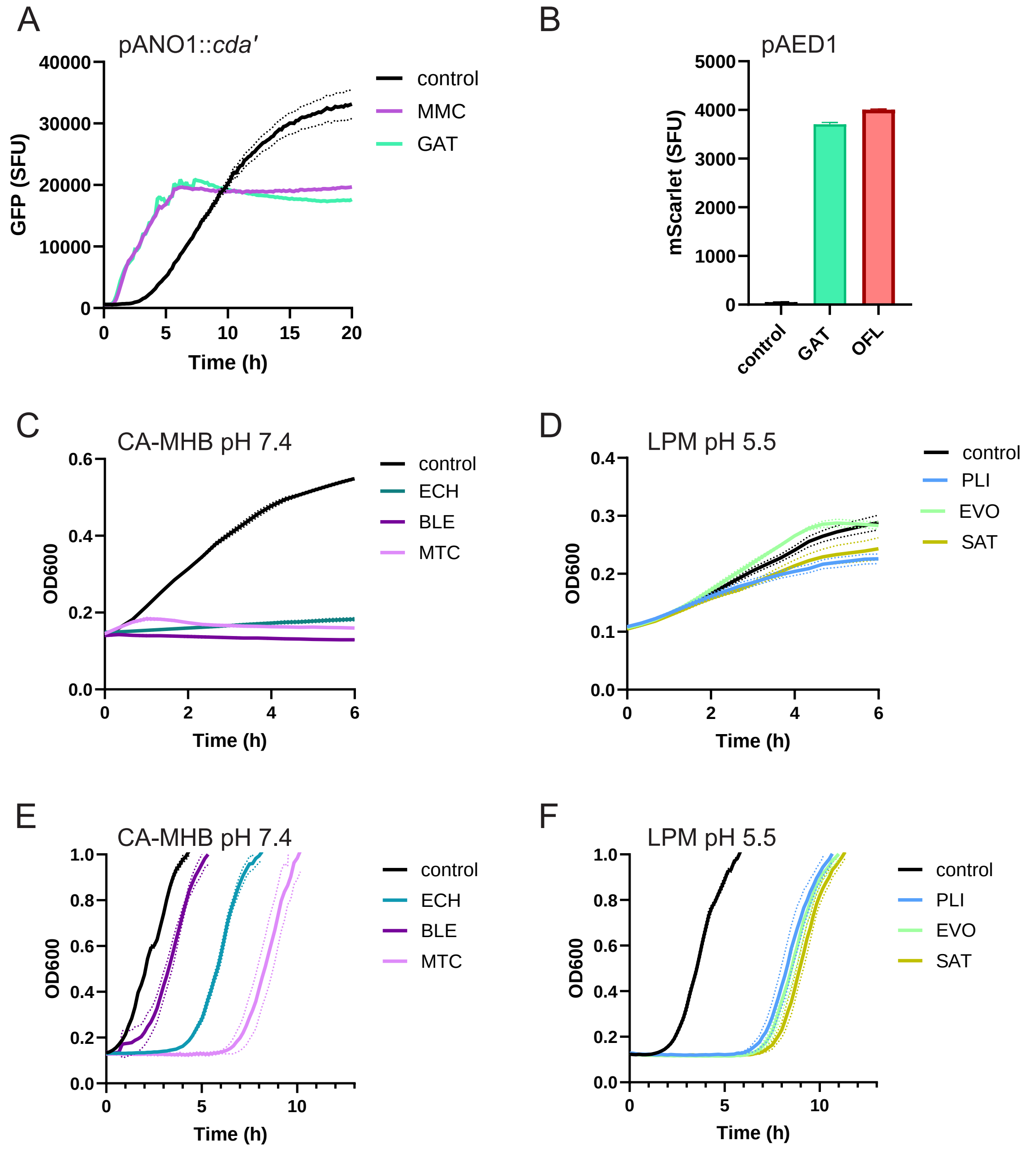


**Figure S5. Activity of anti-cancer compounds on UPEC CFT073**

**A, B.** Validation of fluorescent reporters for the SOS response.
**A.** Cultures bearing the pANO1::*cda'* plasmid were treated with 0.1 µM gatifloxacin (GAT) and 0.5 µM mitomycin C (MMC) or incubated without any drug (control) in a plate reader for 20 h. GFP fluorescence and $OD_{600}$ were recorded every 10 min. Data for the drug-free control represent the means of nine technical replicates ± SEM.
**B.** Cultures bearing the pAED1 plasmid were treated with 0.3 µM GAT and ofloxacin (OFL) or incubated drug-free (control). mScarlet-I fluorescence and $OD_{600}$ were recorded after 6h of incubation. Data represent the means of four technical replicates ± SEM.
**A, B.** Specific Fluorescence Units (SFU) were calculated by normalizing arbitrary fluorescence units to cell density (AU/$OD_{600}$).
**C, D.** Effect of anti-cancer agents on bacterial growth. $OD_{600}$ was measured for cultures bearing pAED1. Bacteria were grown either in 1:4 CA-MHB (pH 7.4) and treated with 20 µM echinomycin (ECH), bleomycin (BLE), and mitomycin C (MMC) (C), or in LPM (pH 5.5) and treated with plicamycin (PLI), evofosfamide (EVO), and satraplatin (SAT) (D).
**E, F.** Effect of anti-cancer agents on regrowth following treatment of stationary-phase cultures. Bacteria bearing pAED1 were cultivated in the same media as in panels C and D for 24 h, then treated with the same compounds for an additional 24 h. Bacteria were collected by centrifugation, washed, resuspended, and diluted 1:7 in CA-MHB. $OD_{600}$ of the regrowing cultures was measured.
**C-F.** Values are presented as means ± SEM for n = 3.

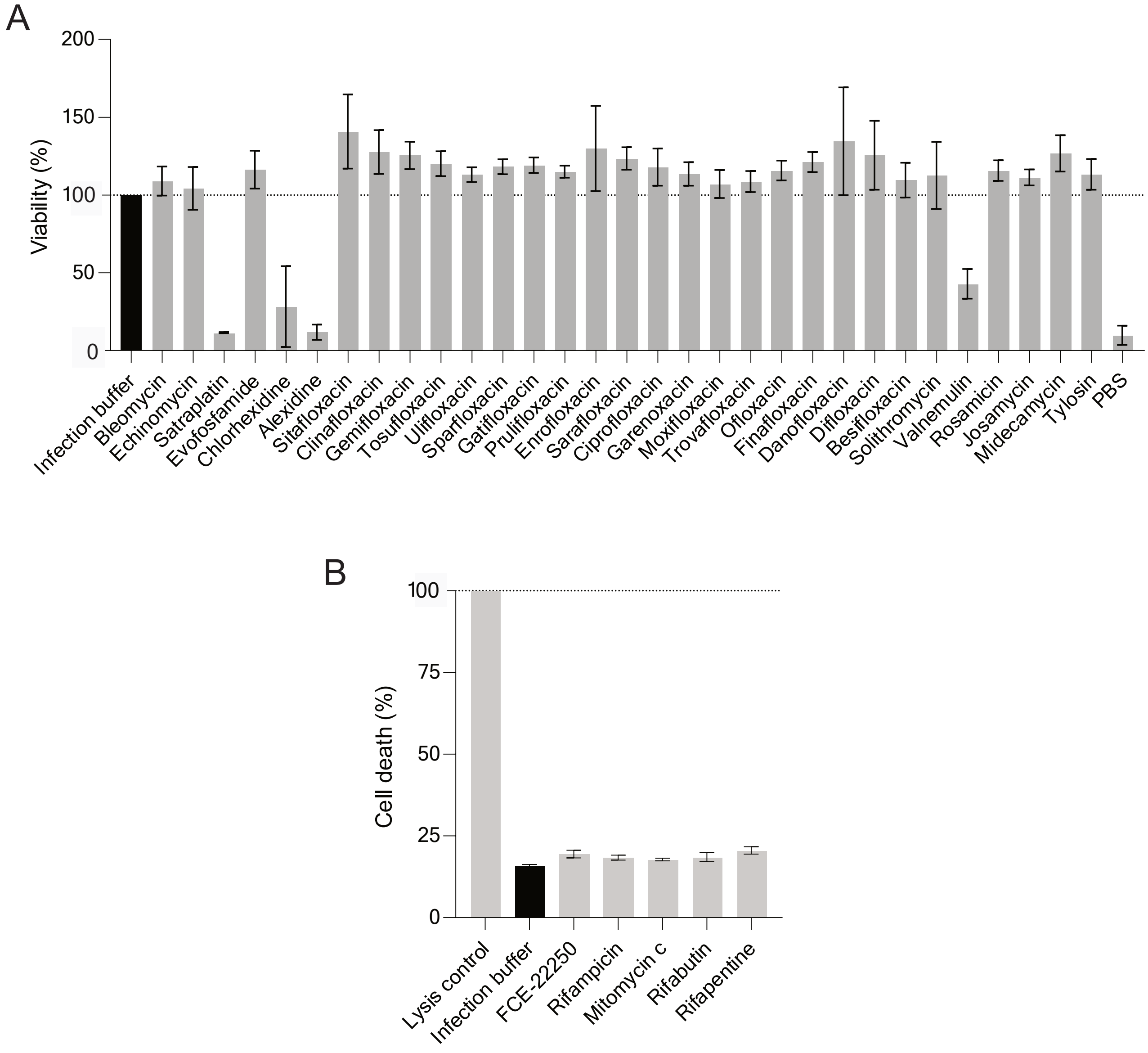


**Figure S6. Impact of hit compounds on TC7 cell viability.**

**A.** *Evaluation of toxicity for non-colored compounds using the MTS assay*. TC7 cells were incubated for 12 h with each compound at its minimum inhibitory concentration (MIC) in the absence of bacteria. CellTiter 96 AQueous One Solution Reagent (Promega) was added to cells incubated with infection buffer. Absorbance was measured at 490 nm after 2 h, and values were normalized to the control wells treated with infection buffer only. The experiment was performed twice, with each condition tested in duplicate. Error bars indicate the standard deviation (SD).

**B.** *Evaluation of toxicity for compounds that absorb light at 490 nm using the LDH release assay.* Cell viability was assessed using the CytoTox-ONE Homogeneous Membrane Integrity Assay kit (Promega) to measure lactate dehydrogenase (LDH) release. Cells in the positive control wells were lysed using lysis solution. Supernatants from each well were incubated with CytoTox 96 Reagent for 10 min, and fluorescence was measured at 560 nm excitation and 590 nm emission. Values were normalized to total lysis controls (representing 100% cell death) and are expressed as means ± SD for n=2.
